# Wakefulness fluctuations elicit behavioural and neural reconfiguration of awareness

**DOI:** 10.1101/155705

**Authors:** Valdas Noreika, Andrés Canales-Johnson, Amy Johnson, Aurina Arnatkevičiūtė, Justin Koh, Srivas Chennu, Tristan A. Bekinschtein

## Abstract

Mapping the reports of awareness and its neural underpinnings is instrumental to understand the limits of human perception. The capacity to become aware of objects in the world may be studied by suppressing faint target stimuli with strong masking stimuli, or – alternatively – by manipulating the level of wakefulness from full alertness to mild drowsiness. By combining these two approaches, we studied how perceptual awareness is modulated by decreasing wakefulness. We found dynamic changes in behavioural and neural signatures of conscious access in humans between awake and drowsy states. Behaviourally, we show a decrease in the steepness of the psychophysical function for conscious access in drowsy trials. Neural mapping showed delayed processing of target-mask interaction as the consciousness transition progressed, suggesting that the brain resolution of conscious access shifts from early sensory/perceptual to decision-making stages of processing. Once the goal to report the awareness of a target is set, the system behaviourally adapts to rapid changes in wakefulness, revealing the flexibility of the neural signatures of conscious access, and its suppression, to maintain performance.

**Significance statement:** Maintaining full alertness for long periods of time in attentionally demanding situations is challenging and may lead to a decrease in performance. We show the effect of wakefulness fluctuations on behaviour and brain dynamics that humans use to maintain performance. We reveal the neural strategies we have to cope with drowsiness by shifting the weights to more flexible brain processes and relaxing the precision of the decisions we take.

## INTRODUCTION

In the past decade, two independent research programs have led the study of the neural correlates of consciousness (NCC): the first focuses on the neural mechanisms of conscious awareness and access(1–5), while the other one seeks to explain the neural basis of the states of consciousness(6–11). Conscious access, i.e. reportable awareness of external input, is typically studied by presenting target stimuli around a sensory/perceptual threshold value, enabling the contrast between trials of presence versus absence of reported awareness. A paradigmatic example of this approach is the presentation of targets with variable intensities of noise, rendering them inaccessible in a subset of trials(12, 13).

Proposed neural correlates of conscious access could be grouped into relatively ‘early’ processes such as recurrent processing within sensory cortices and sensory-central interactions (~100-250 ms post-stimulus)(5, 14), and relatively ‘late’ contribution of central executive processes (~250-400 ms post-stimulus)(15, 16). While a theoretical disagreement persists as to whether the early candidates for NCC reflect a ‘pre-conscious’(17) or a ‘phenomenally conscious’(8, 18) stage of processing, a general consensus is that reportable conscious access is associated with the late executive NCC, although a more complex picture may be more informative(19). The Global Neuronal Workspace model of consciousness(2, 3) interprets the late NCC as a stage of processing where specific information of sensory input is amplified and re-encoded in the prefrontal–parietal network(15, 20, 21). As such, conscious access is characterized, both behaviourally and neurophysiologically, by a sigmoidal relationship between performance/brain markers and degree of masking(16, 22, 23).

Instead of contrasting trials that differ in the awareness of particular stimuli, the second group of NCC studies aims to contrast different states of consciousness, such as awake vs. sedated(24–26), awake vs. anaesthetized(27, 28), awake vs. asleep(11, 29, 30), dreaming vs. dreamless sleep(10, 31), or minimally conscious state vs. unresponsive wakefulness syndrome(32, 33). Typically, the states of consciousness are characterized by relatively stable neural processes that range between several hundreds of milliseconds to minutes and can be measured by indices of neural dynamics (power and oscillatory) and information measures(26, 29, 33–36). From the perspective of the conscious access program of research, the neural signatures of state are typically regarded as an enabling factor determining the capacity for conscious access(2, 37, 38). That is, the neural markers of conscious states are seen as necessary to enable conscious access, but their proposed role is limited to arousal modulation, and their likely influence on conscious access mechanism has not been considered, nor included dynamically in integrative models of consciousness(7, 39). In contrast to this mainstream view, we propose that arousal is more than an enabling factor and may in fact differentially modulate the neural processes underlying conscious access.

In this study we integrated the conscious access and the consciousness state programs by studying behavioural and neural dynamics of auditory backward-masked targets in the transition from awake to sleep(40). In particular, we modulated awareness and wakefulness along two dimensions: **(i)** conscious access was externally manipulated by masking the auditory targets, and **(ii)** the overall level of wakefulness was endogenously decreased by facilitating natural transition from awake to drowsy states. This experimental paradigm reduced wakefulness and yielded a considerable proportion of drowsy yet responsive trials. This way, we were able to study the interaction between conscious access (“aware” vs. “unaware” trials) and conscious states (“awake” vs. “drowsy” trials). Based on the premises that **(i)** threshold and slope of psychophysical functions depend on the noise in the system(41–44), and that **(ii)** decreased wakefulness increases noise in the sensory-perceptual and fronto-parietal networks(45–47), we hypothesized that the threshold of auditory awareness will increase and the slope will decrease as a function of decreasing wakefulness. Furthermore, we expected that corresponding sensory and executive electrophysiological markers of conscious access will show delayed dynamics in a drowsy state of consciousness. Indeed, our findings confirm that both the behavioural and electrophysiological signatures of conscious access are altered by changing the state of wakefulness.

## RESULTS

To investigate the neural correlates of conscious access in the transition from wakefulness to sleep, human participants (N=31) carried out an auditory detection task where they judged the presence or absence of a target tone followed by varying silent gaps before a white noise mask was presented. The participants were encouraged to relax in a reclining chair during the 3 hours of the task to drift into drowsy states (but awoken if unresponsive). On test trials, a target sound was presented and then followed by a masking noise after one of 11 intervals of silence centred on an individualised target detection threshold (see Fig 1A). Participants were instructed to report if they have heard a target sound or not after a response cue. High-density electroencephalography (EEG) measurements were carried out throughout the experiment. To assess the fluctuating level of wakefulness at a single trial level, two EEG measures were derived from the spontaneous pre-stimulus neural data. First, the ratio of theta/alpha (θ/α: 4-7 Hz/8-12 Hz) spectral power over a 2 sec pre-stimulus period was used for behavioural analyses (see Fig 1A, S1 and Online Methods); and second, Hori stages of sleepiness over a 4 sec pre-trial period were assessed for EEG analyses(39, 48, 49). As a complementary behavioural measure of wakefulness, reaction times (RT) locked to the onset of response cue were also calculated.

**Figure 1.**
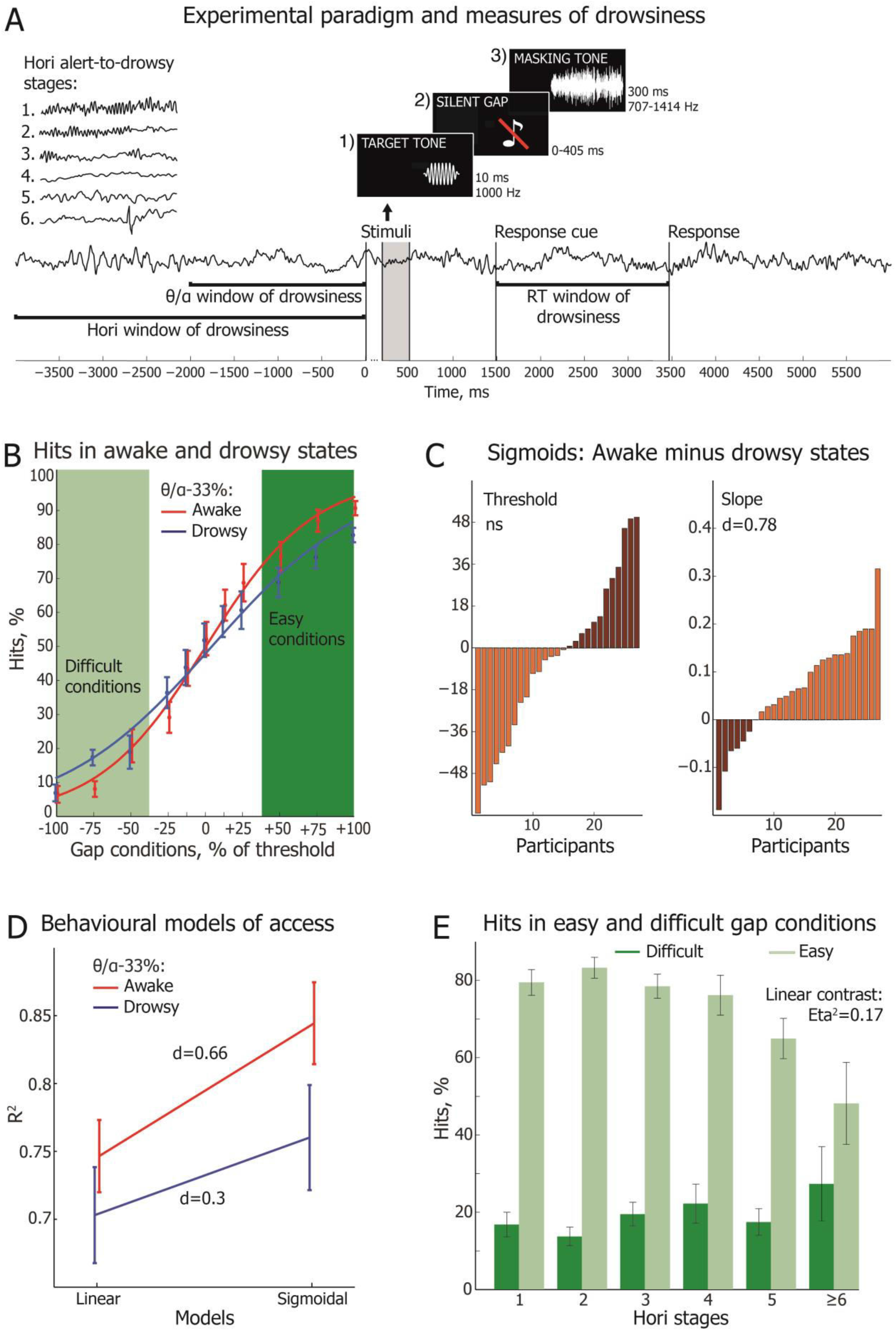
Experimental design, slope and threshold of conscious access in the EEG-defined awake and drowsy states. Once participants became drowsy, they showed a decrease in the slope of the psychophysical function with higher fit to linear than sigmoidal models and a decreasing difference between the easy and difficult conditions in the proportion of hits. **(A)** Trial structure in a time window of a single EEG epoch (−4000 ms to 6000 ms) in relation to the onset of the target tone. The 300 ms mask (grey shaded bar) was followed by a response cue after 1000 ms. Responses beyond 6000 ms from the onset of target were not considered and the trial was regarded as unresponsive. Green lines indicate time windows of the three measures of wakefulness: −4000 ms to 0 ms period was used for scoring Hori stages of sleep onset (in this example, Hori Stage 5 is shown in the right occipital channel); −2000 ms to 0 ms time window was used to calculate EEG spectral power ratio between theta (4-7 Hz) and alpha (8-12 Hz) frequency bands. **(A inset)** EEG examples and brief definitions of 6 Hori stages of sleep onset progression from the relaxed wakefulness to the appearance of vertex sharp waves. **(B)** Sigmoidal functions fitted to awake (red) and drowsy (blue) hits, following θ/α-33% top/bottom split of the states of consciousness. The trials were first averaged within each participant and then across all participants. The error bars indicate the standard error of mean (SEM), calculated across all participants. **(C)** Threshold and slope estimates in the drowsy state subtracted from those in the awake state split by θ/α-33%. Each individual participant (N=27) is represented in one bar and sorted from the largest increase of threshold (light brown) to the largest decrease (dark brown) of threshold in drowsy state (left); the same for slope. **(D)** Goodness of fit (coefficient of determination, R^2^) of linear and sigmoidal models of the proportion of hits as a function of gap conditions in the awake and drowsy trials as defined by θ/α-33%. The error bars indicate the SEM. **(E)** Proportion of hits in the right tail of sigmoid (easy gap conditions +50%, +75%, +100%; light green) and in the left tail of sigmoid (difficult gap conditions −100%, −75%, −50%; dark green) across Hori stages. A difference between the easy and difficult conditions decreases with decreasing wakefulness. The error bars indicate the SEM.

### The transition of consciousness modulates behavioural markers of conscious access

To compare distribution of a hit rate as a function of conscious state, a sigmoid function was fitted across 11 gap conditions for each participant separately for the θ/α- and RT-defined awake and drowsy trials, yielding individual threshold and slope estimates of the psychophysical function for both states of consciousness. While there was no significant difference in the threshold of conscious access between awake and drowsy trials at the group level, we observed a robust decrease of sigmoid slope in drowsy state of consciousness in the group and in the majority of the participants (see Fig 1B-C, S2 and Table S1). Notably, this decrease in slope with decreased wakefulness was robust and convergent when assessed with the four alternative splits of data defined by θ/α and RT measures of wakefulness (θ/α-33%, θ/α-50%, RT-33%, and RT-50%; see Online Methods). In all cases, the decreasing slope of sigmoid function indicated that conscious access becomes more “gradual” in the transition from wakefulness to sleep.

To test formally if the dynamics of conscious access differs between states of consciousness, we analysed how well the behavioural data fit a sigmoidal versus a linear model, comparing the coefficient of determination R^2^ of both models in the θ/α- and RT-defined awake and drowsy states (see Fig 1D). A significant Models x States interaction was observed (θ/α-33%: F(1,26)=13.82, p=0.0009, see Fig 1D; RT-33%: F(1,25)=33.34, p<0.000005, see Fig S2C). Even though sigmoidal models had a higher R^2^ than linear models in both states of consciousness, the difference between the models was lower in drowsy (θ/α-33%: M=0.057, SEM=0.015; RT-33%: M=0.037, SEM=0.01) than in awake (θ/α-33%: M=0.098, SEM=0.014, t(26)=3.72, p=0.001, d=0.54; RT-33%: M=0.114, SEM=0.015, t(25)=5.77, p<0.000005, d=5.97). These findings point to a change of conscious access dynamics as alertness wanes. The slope of conscious access decreases in drowsy states of consciousness and the distinction between “aware” and “unaware” decision becomes less dichotomous.

Hypothetically, the decrease of slope in trials split by EEG θ/α power ratio may be dependent on neurocognitive processes other than changes in wakefulness. In particular, changes in pre-stimulus alpha power may reflect endogenous fluctuation of attentional sampling and/or sensory gating(50–52). To examine this alternative, we carried out a complementary analysis based on Hori-defined stages of the transition from wakefulness to sleep. For each individual, the percentage of hits in the most difficult gap conditions ‘-100%’, ‘-75%’, ‘-50%’ was subtracted from the percentage of hits in the easiest gap conditions ‘+50%’, +75%’, ‘+100%’, separately for each of the available Hori Stages 1-6. The difference score between the easy and difficult conditions decreased from Hori Stage 1 to Stage 6 (weighted linear term ANOVA: F(1,5)=30.18, p<0.00005, Eta^2^=0.17; see Fig 1E), confirming that participants gave less “aware” responses in the easy trials and more “aware” responses in the difficult trials when the level of wakefulness decreased, suggesting a reduced capability to discern between difficult and easy trials in a drowsy state. This convergent finding corresponds to a decrease of slope observed in the RT- and θ/α-based analyses, confirming a reduced steepness of conscious access in a drowsy state of consciousness.

Interestingly, threshold estimates in the θ/α- and RT-awake state were strongly associated with threshold estimates in drowsy (coefficient r: 0.52-0.77, see Table S1). Likewise, slope estimates correlated positively between awake and drowsy states of consciousness as defined by θ/α (coefficient r: 0.2-0.78, see Table S1). That is, participants who performed well in awake were also more accurate in drowsy state, showing that individual traits of threshold and slope of auditory access may be relatively stable across the two states of consciousness. Contrary to this, threshold and slope were not associated in any of the θ/α- or RT-based data splits of data (see Table S1), suggesting that these two behavioural measures of conscious access are independent. This observation is consistent with the slope decreasing with lower wakefulness while the threshold remains unaffected. Overall, behavioural findings revealed that the distinctive non-linear dynamics of conscious access might be a flexible feature of consciousness, dependent on its state.

### Transition to sleep modulates neural markers of conscious access

To study whether neural markers of conscious access may also be state-dependent, we first determined a sequence of neural processing stages, depicted by event related potentials (ERP), which distinguish aware and unaware trials. We found that hits are associated with increased amplitude of N100 ERP cluster in response to target tone, and decreased amplitude of N100, P200 and P300 ERP clusters in response to mask (see Fig S3 and Supplementary Results). That is, the suppression of conscious access was associated with enhanced amplitude of potentials evoked by the masking noise. To determine which of the identified ERP markers of conscious access are modulated by the state of wakefulness, we further split the data between awake (Hori 1-2 stages) and drowsy (Hori 4-5 stages) trials, and analysed States (awake vs. drowsy) and Access (hits vs. misses) interaction at four stages of neural processing: (1) facilitatory sensory ignition by a target tone (target N100); (2) inhibitory sensory ignition by a noise (mask N100); (3) inhibitory sensory-attentional gating of noise (mask P200); or (4) inhibitory global broadcasting of noise (mask P300).

While the target-evoked N100 cluster showed higher amplitude in the awake than in the drowsy trials, the relationship between hits and misses remained the same in both states of consciousness (see Fig 2A), i.e. a changing brain state did not seem to modulate access NCC at the earliest sensory ignition stage of target-evoked auditory processing. However, a clear Access x State interaction was observed for the mask-evoked N100 cluster, with misses being associated with higher amplitude than hits in the awake, but not in the drowsy state of consciousness (see Fig 2B). That is, sensory processes evoked by a masking noise interfered with conscious access only when wakefulness level was relatively high. A similar state-dependent modulation of conscious access was observed in the analysis of mask-evoked P200 cluster: misses had higher cluster amplitude than hits in the awake but not the drowsy state of consciousness (see Fig 2C and Fig S4). Regarding mask-evoked P300 cluster, misses were associated with higher amplitude than hits in both states of consciousness, with no significant Access x State interaction (see Fig 2D).

**Figure 2.**
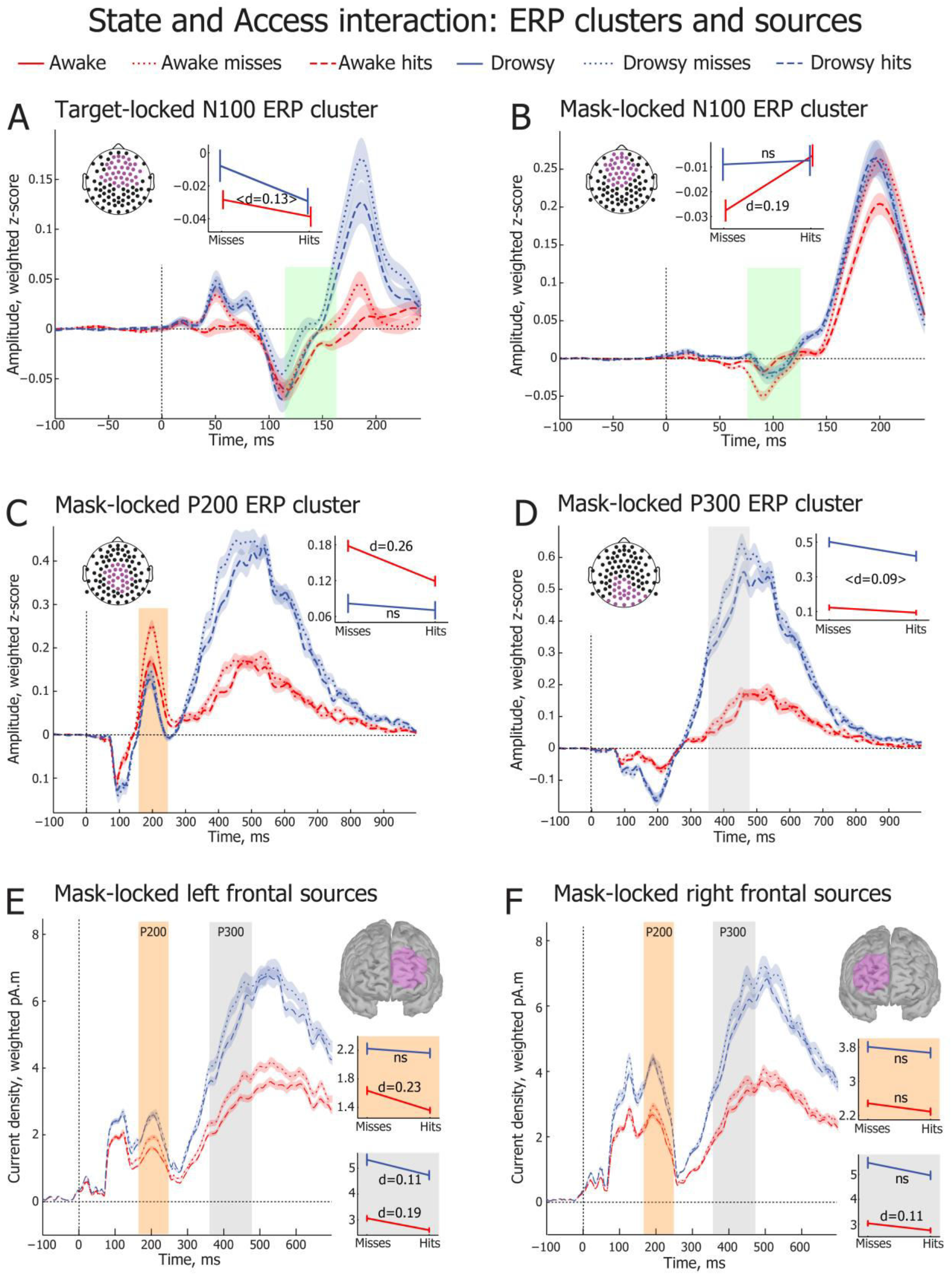
Wakefulness modulates neural signatures of conscious access suppression. **(A-D)** Time courses of N100, P200 and P300 waveforms locked to the onset of target/mask (0 ms). Waveforms are averaged across all cluster electrodes, which are depicted in violet in the electrode montage maps. Shaded vertical bars indicate cluster time windows used to calculate the mean amplitude of N100 (green), P200 (orange) and P300 (grey) clusters. Individual trials are averaged separately for the four conditions: awake misses, awake hits, drowsy misses, and drowsy hits. Insets depict interaction plots of State (awake, drowsy) and Access (miss, hit) factors for the target-locked N100 and mask-locked N100, P200, and P300 ERP clusters. **(A)** Target-evoked N100 cluster revealed significant main effects of State (F(_1,1272_)=4.48, p=0.035) and Access (F(_1,1272_)=5.03, p=0.025), whereas the States x Access interaction was not significant (F(_1,1272_)=0.63, p=0.43). With awake and drowsy trials grouped together, hits were associated with higher target-evoked N100 amplitude than misses (t(_1274_)=2.24, p=0.025, d=0.13). **(B)** Mask-evoked N100 cluster revealed significant main effect of Access (F(1,3168)=5.24, p=0.022), and a weak State x Access interaction (F(1,3168)=3.887, p=0.049), whereas the main effect of State was not significant (F(1,3168)=2.55, p=0.11). Misses had higher amplitude than hits in the awake trials (t(1584)=3.81, p=0.00015, d=0.19), but not in the drowsy trials (t(1584)=0.19, p=0.85, d=0.01). **(C)** Mask-evoked P200 cluster revealed significant main effects of State (F(1,3168)=39.45, p<0.000005) and Access (F(1,3168)=9.3, p=0.0023) and a weak State x Access interaction (F(1,3168)=4.28, p=0.039). Misses had higher P200 amplitude than hits in the awake trials (t(1584)=5.18, p<0.000005, d=0.26), but not in the drowsy trials (t(1584)=0.56, p=0.57, d=0.03). **(D)** Mask-evoked P300 cluster revealed significant main effects of State (F(1,3168)=294.75, p<10^-8^) and Access (F(1,3168)=7.43, p=0.0064), whereas the State x Access interaction was not significant (F(1,3168)=1.67, p=0.197). With awake and drowsy trials grouped together, misses were associated with higher mask-evoked P300 amplitude than hits (t(3161)=2.61, p=0.0092, d=0.09). **(E-F)** Weighted minimum norm estimation (wMNE) of single trial frontal sources (violet) averaged across all four conditions (awake misses, awake hits, drowsy misses, and drowsy hits) in the P200 and P300 ERP cluster time windows. Waveforms indicate time courses of source activations, whereas statistical plots depict planned pairwise comparisons of frontal source activations between hits and misses in awake and drowsy states in P200 (orange) and P300 (grey) time windows. Error bars indicate SEM. **(E)** In the P200 time window, awake misses had higher amplitude than awake hits in the left frontal (t(1433.7)=4.32, p=0.000017, d=0.23) source. **(F)** In the P300 time window, awake misses had higher amplitude than awake hits in the left frontal (t(1507.3)=3.72, p=0.00021, d=0.19), and the right frontal (t(1570)=2.16, p=0.031, d=0.11) sources. Drowsy misses showed a higher amplitude than drowsy hits in the left frontal source (t(1514.7)=2.22, p=0.027, d=0.11).

These findings indicate that while conscious access was initially associated with higher amplitude of target-evoked N100 cluster in both awake and drowsy states, the final resolution of target-mask interaction seemed delayed in drowsy state. In particular, awake misses had higher mask-locked N100 and P200 neural activity than awake hits, whereas no reliable differences were observed between drowsy misses and hits in the same N100 and P200 time windows. Masking of conscious access in decreased wakefulness efficiently supressed neural markers of early stages of processing (N100 and P200), but did not sufficiently affect central resources for conscious awareness of targets at later stages of processing (P300). These ERP findings may indicate that the mechanisms of conscious access and its inhibition are not hardwired; instead, they are state-dependent and flexible. Mask-evoked N100 and P200 clusters with relatively low amplitude may not contribute to the inhibition of conscious access in drowsy trials, while a high-amplitude P300 cluster differentiated aware and unaware trials in both levels of wakefulness. Conscious access is resolved at the early processing stages (N100/P200) during normal level of wakefulness (Hori stages 1-2), while during drowsy state marked by EEG flattening and theta waves (Hori stage 4-5), awareness is resolved only in the late stages of the neural processing (P300) (see Fig 2B-D).

We expected the suppression of auditory conscious access to involve wide broadcasting of masking noise in the temporal and fronto-parietal nodes of processing, sparing no serial resources for the maintenance of a target tone(2, 15). Following previous research into fronto-parietal functioning during drowsy states(46, 47), we further hypothesized that wakefulness would differentially modulate fronto-parietal network in these critical stages of mask processing. To address these hypotheses, we carried out exploratory single-trial source analysis by averaging activation of temporal and fronto-parietal sources locked to the onset of mask (see Fig S5 A). Afterwards, the trials were split between Access (hits vs. miss) and State (awake vs. drowsy) conditions. The main effect of Access was observed in P200 (F(1,3168)=5.47, p=0.019) and P300 (F(1,3168)=5.1, p=0.024), but not in N100 (F(1,3168)=0.16, p=0.69) time windows. Similar to the sensor-level statistics, analysis in the source space showed a weak State x Access interaction for the P200 cluster (F(1,3168)=3.99, p=0.046), with misses having higher amplitude than hits in the Hori-awake (t(1563.8)=3.98, p=0.00007, d=0.2), but not in the Hori-drowsy (t(1584)=0.2, p=0.84, d=0.01) states of consciousness. Contrary to this, the States x Access interaction was not noticeable in the P300 time window (F(1,3168)=0.00, p=0.98), with misses having marginally higher P300 amplitude than hits in both awake and drowsy trials (t(3170)=2.09, p=0.037, d=0.07). These findings may confirm our earlier observation at the sensor level that the resolution of conscious access is delayed in drowsy state to the final P300 stage of mask processing, whereas an earlier target-mask interaction may anticipate if a target will be reported in awake trials.

To identify which of the 6 predefined regions of interest (left and right frontal, parietal and temporal, see Fig S5 A) show an expected increase of amplitude in misses compared to hits, we carried out planned pairwise comparisons for each source and state of consciousness. In the P200 time window, the mask evoked higher activation of the bilateral temporal and the left parietal and frontal sources in the Hori-awake trials, whereas no differences between hits and misses were observed in Hori-drowsy trials (see Fig 2 E-F and Fig S5). That is, suppression of conscious access in the P200 time window showed a correlate of neuronal processing in both perceptual and central networks (temporal and fronto-parietal) in the awake but not in the drowsy state. In the P300 time window, a higher amplitude in misses than hits was observed in the left parietal (see Fig S5 B) and bilateral frontal sources in the awake state, but only in the left frontal source in the drowsy state of consciousness (see Fig 2 E-F). Even though the effects were allegedly small (Cohen’s d<0.20), these findings suggest that in addition to the temporal restriction of neural markers (late for drowsy), the neural network underlying reported awareness shrinks during transition to sleep, despite the higher general activation, indexed by increased P300 amplitude in drowsy states (see Fig S4).

### Transition to sleep modulates sensory-central gating

Given that temporal proximity of the target and mask sounds largely determines if target awareness will be reported (see Fig 1B), we were interested whether the temporal proximity of target and mask differentially modulates ERP clusters in the awake and drowsy states. We hypothesized that electrophysiological signature of this proximity will coincide with the first stages of neural processing leading to conscious access. In particular, neuronal responses to suppressive masking noise should change as a function of a delay from the preceding target tone. That is, mask-evoked ERP responses should have higher amplitude in the most difficult gap conditions when conscious access is suppressed, and smaller amplitude in the easiest gap conditions when target tones escape suppression. More specifically, we hypothesized that the delay between target and mask modulates mask-evoked responses in N100 or P200 time windows when sensory information becomes available for central processing, i.e. during sensory- or sensory-central gating(53). Contrary to this, P300 as a marker of the final stages of processing of conscious access was not expected to expose relatively early sensory-central gating as it reflects central processes rather than perceptual processing of stimuli(54, 55). To test these hypotheses, we fitted a linear function to the mean amplitude of ERP responses in the N100, P200, and P300 cluster time windows across 11 gap conditions, separately for each electrode, and then analysed topographical heat maps of slope values (see Fig 3).

**Figure 3.**
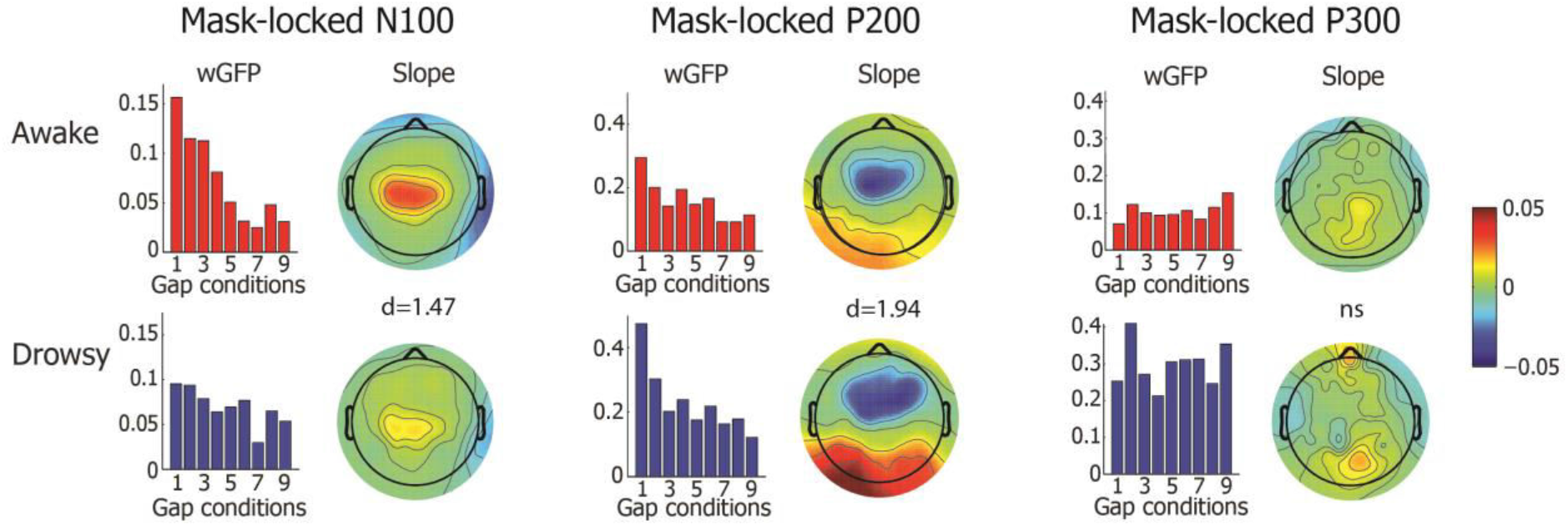
Target-mask interaction revealed by a linear fit of ERP responses across gap conditions. Linear fit of the mask-locked N100 (left), P200 (middle) and P300 (right) ERP cluster amplitudes across target-to-mask silent gap conditions. Bar plots depict weighted global field power (wGFP) in a time window of respective cluster across 9 equally spaced gap conditions. Conditions ‘-12.5%’ and ‘+12.5%’ are not presented here, as they were additional intermediate intervals. Scalp maps represent slope values of a linear function, which was fitted to the mean amplitude of respective ERP clusters across 11 gap conditions. The slope was first calculated for each electrode and the obtained values were interpolated in the heat maps. The top line represents fit of the awake trials (Hori 1-2), whereas the bottom line depicts fit of the drowsy trials (Hori 4-5).

N100 and P200 maps showed relatively high slope values, indicating that amplitude of mask-evoked potentials reflect temporal proximity of target-masking sounds in these time windows. Contrary to this, slope values were very low in P300 time window (see Fig 3). Furthermore, we observed that sensory-central gating in N100 and P200 time windows is state-dependent. Repeated measures ANOVA of 92 slope values representing each electrode with the State (Awake, Drowsy) and Component (N100, P200) as independent factors revealed a significant main effect of the ERP components (F(1,91)=41.62, p<0.000005), the States (F(1,91)=10.47, p=0.0017), and a strong Components x States interaction (F(1,91)=313.35, p<0.000005). In particular, there was a steeper decrease of N100 amplitude across the gap conditions in awake compared to drowsy state (t(91)=6.99, p<0.000005, d=1.47), whereas P200 slope was steeper in drowsy than awake state (t(91)=9.25, p<0.000005, d=1.94) of consciousness (see Fig 3 and Fig S6). Furthermore, a difference between ERP slope in N100 and P200 cluster windows was higher in the drowsy state (t(91)=9.35, p<0.000005, d=1.96) than in awake state (t(91)=2.21, p=0.029, d=0.46).

These findings indicate that temporal proximity of target and masking sounds, a key parameter for conscious access and its suppression, is processed at a relatively early N100 time window in the awake state of consciousness, and that this stage of processing is delayed to the P200 time window in drowsy state. That is, sensory-central gating becomes more sluggish in the transition from wakefulness to sleep, and probably more neural resources are required to determine the presence of the target followed by the mask during drowsy states.

## DISCUSSION

Our findings demonstrate that behavioural and neural markers of conscious access are conscious state-dependent. In particular, we observed a more gradual interaction between the masking intensity and the rate of target detection when participants became drowsy compared to more abrupt conscious access dynamics in awake state. A more gradual build-up of evidence for conscious access(56) in decreased wakefulness may depend on the increased variance in the underlying neural processes that could produce a shallower response slope(57). At the behavioural level, the increase of a hit rate in the most difficult trials with its simultaneous decrease in the easiest trials when the threshold point remains constant suggest detection instability in a proportion of drowsy trials as compared to fully awake. This finding implies that a steep detection slope(16) may not be a fundamental property of conscious access, but instead it can change as a function of wakefulness microstate before a stimulus appears. Thus, the opposing gradual(58) vs. non-gradual(59) models of conscious access may not be mutually exclusive; instead, some aspects of their mismatch can be reconciled by adding a factor of a varying level of wakefulness, which may modulate attention and awareness(7).

Wakefulness thus seems to provide a distinctive behavioural distortion of consciousness, which takes place during a period of rapid fronto-parietal reorganization(46, 47, 60), as further indicated by the increased P300 amplitude in the present study. It has been observed previously that neurological patients with prefrontal lesions have an increased threshold yet a stable slope of conscious access(55). Contrary to this, and to our expectation, transition to drowsy states uncovered a unique case of a decreasing slope yet spared threshold of conscious access, which differs orthogonally from the earlier neurological observations(55). The possibility that either a slope or a threshold can be selectively modulated by a change of a brain state suggests that these two behavioural markers of consciousness are partially independent, further confirmed by the absence of their correlation in the present study.

At the electrophysiological level of analysis, we identified a sequence of evoked potentials that distinguished hits and misses across different stages of a complex target-mask interaction. In particular, hits were associated with higher amplitude of target-evoked potential in the N100 time window (peaking at 150 ms), replicating earlier ERP studies of auditory masking(61–63). Given that the N100 marker of auditory detection is generated primarily in the auditory cortex(62), our finding provides support to the ‘early’ sensory NCC accounts of conscious access(5, 14) for auditory signals. In the later stages of target-mask interaction, the initial representation of target was suppressed by mask-evoked N100, P200, and P300 potentials. Thus, in addition to the previous report that misses have higher P200 amplitude(64), we show that mask evoked activity interferes with conscious access at each of the perceptual (N100), perceptual-central (P200), and central (P300) stages of information processing.

After splitting the EEG data between awake and drowsy trials, we found evidence that EEG markers of conscious access are also state-dependent. In particular, the graduality of mask-evoked N100 response across 11 gap conditions in the awake state was shifted to the P200 time window when participants became drowsy. This finding suggests that the drowsy brain requires the involvement of further central resources for the distinction between two serially presented stimuli. Arguably, delayed processing of the target-mask interaction may also contribute to the decreasing sigmoidal slope of conscious access in drowsy state.

The sluggishness of EEG response in decreased wakefulness was also observed when interaction between conscious access (hits, misses) and state (awake, drowsy) was studied in the sensor as well as the source neural spaces. While target-evoked N100 had higher amplitude in hits than in misses in both states of consciousness, mask-evoked suppression of conscious access occurred only in P300 time window in drowsy state compared to significant effects observed in mask-evoked N100, P200 and P300 time windows in the awake state of consciousness. Conscious access was thus associated with a decreased response to mask, i.e. a higher resistance to the interfering sound, in all three stages of processing in the awake but not in the drowsy state of consciousness. Notably, mask evoked N100 and P200 potentials had relatively low amplitude compared to P300 potentials in drowsy states (see Fig S4), and likewise neuronal processes that distinguished hits and misses were resolved in the time window of the largest ERP response. These observations seem to point to the multiple realization of the NCC(37) in different states within the same participants. That is, the neural markers of consciousness and its suppression appear to be flexible and dependent on the overall state of the brain, and an NCC observed in one state may not hold for another state of consciousness. Alternatively, a possibility cannot be ruled out that the subjective quality of auditory awareness differed between awake and drowsy states, which may have involved correspondingly different NCC(65).

Despite state-dependent differences in the association between conscious access and N100 / P200 clusters, the final P300 stage of the neuronal cascade towards conscious access distinguished hits and misses in both states of consciousness. Source modelling of P300 revealed a fronto-parietal activation that likely reflects global broadcasting of suppressive masking noise(2). While three of the four P300 sources distinguished aware and unaware trials in awake trials – the left and right frontal and the left parietal cortex – only the left frontal source showed a higher activation in misses than hits in drowsy trials. Interestingly, patients with lesions in the left compared to the right prefrontal cortex also show stronger suppression of conscious access(55). The left frontal component of the global broadcasting network appears to be critical for the conscious access, and is the last one to survive a state of decreasing level of wakefulness while participants remain responsive.

Previous attempt to study conscious access in different states relied on pharmacological intervention(66). Transition to sleep provides a fruitful naturally occurring alteration of the state of consciousness with a preserved capacity to respond, which can be utilized to study the modulation of conscious access by a changing brain-state. Such natural fluctuation of the level of wakefulness may also occur unintentionally and go undetected(36), which – if left uncontrolled – may hinder attempts to narrow down neural processes associated with particular contents of consciousness (19).

Our study indicates that configuration of neural processing stages underlying cognitive performance may fluctuate within the same participant at a rapid rate within a single session of experiment. Yet, despite decreased level of wakefulness, responsive participants can maintain their goal-set to report auditory awareness, demonstrating flexibility of human brain to adapt to increasing levels of both exogenous (masking) and endogenous (arousal) noise. We have previously shown that individuals maintain capacity to categorize semantic categories of words even during NREM sleep(30), although it remained uncertain if they were aware of presented stimuli. In the present study, participants were conscious in both awake and drowsy trials, revealing that different modes of evidence accumulation, such as a decreased behavioural slope and ERP reconfiguration with lower wakefulness, can underlie conscious access in different states of consciousness. We thus conclude that a comprehensive model of conscious access should include an independent factor of the state of wakefulness, and that access NCC should not be studied in isolation of state NCC(67).

## MATERIALS AND METHODS

### Participants

56 participants (23 male; mean age 26.9; age range 19-39) signed informed consent and took part in the study. Inclusion criteria were being 18 to 40 years old, having no history of hearing impairment or injury and no neurological or psychiatric disorders, and being right handed, which was assessed with the Edinburgh Handedness Scale(68). In order to recruit individuals who are likely to become drowsy and fall asleep in a suitable setting during daytime, potential participants were screened with the Epworth Sleepiness Scale(69), aiming to have a minimum daytime sleepiness score of 7. Given the difficulty in carrying out a very demanding auditory detection task for a long period of time (2 hours), a large proportion of participants failed to maintain a sufficient level of accuracy throughout the session, e.g. their false alarm rate was more than 50% in catch trials, and/or sigmoid function could not be fitted to their responses, especially in a drowsy state of consciousness. These participants were excluded from the analyses reported here, and the final sample consisted of 31 participants (9 male; mean age 27.4; age range 20-39). Notably, the excluded participants reached a deeper level of wakefulness as measured by the intra-individual mode of Hori scores of sleepiness (see Section 2.4.1; inter-individual Mode=5; Range=4) than the remaining participants (inter-individual Mode=2; Range=4; Mann-Whitney U: Z=2.23; p=0.026). Thus, it is likely that the failure to maintain a sufficiently high level of response accuracy was due to a deep level of sleepiness. Interestingly, the excluded participants were also more likely to be male (Pearson χ^2^=4.16, p=0.041), who are generally known to have shorter sleep latency than female adults(70).

The experimental protocol was approved by the Cambridge Psychology Research Ethics Committee (2012.15), and the study was carried out in accordance with the Declaration of Helsinki. Participants were recruited through the electronic volunteer database of the MRC Cognition and Brain Sciences Unit by posting an online advertisement about the study. They received £30 for taking part in the study.

### Experimental design

After signing informed consent, participants filled in the handedness and sleepiness questionnaires, and were seated in a shielded chamber of the EEG lab. While placing the EEG net, participants were instructed that they will have to attend to sounds and report if they have heard a target sound, i.e. a tiny beep, which may or may not occur before a masking sound of noise. Notably, this was not a forced-choice task and participants were not asked to guess if there was a target or not; instead, they were instructed to report if they could hear the target (an “aware” or a “hit” trial) or not (an “unaware” or a “miss” trial). Once the EEG net was applied, participants laid in a fold-back chair in a relaxed position with eyes closed in a dark room. First, a behavioural staircase experiment was carried out in order to estimate individual threshold, i.e. a minimal duration of a silent gap between the target and masking sounds required for the detection of targets. Next, the main EEG experiment was carried out using the individual threshold that was estimated during the behavioural task. Auditory stimuli were presented binaurally using Etymotics ER-3A earphones, and the volume of sound was individually adjusted at a comfortable level.

### Behavioural staircase procedure

During behavioural experiment, a series of masked target sounds were played, and participants responded each time if they could hear the target or not (for the trial structure, see Fig 1A). Each trial began with a target stimulus (10 ms; 1000 Hz; fade-in and fade-out: 2.5 ms each; attenuation: −24 Db), followed by a masking noise (300 ms; fade-in and fade-out: 5 ms; attenuation: 0 Db). The frequency range of the masking sound (*FM*) was octave wide around the frequency of the target sound (*FT*) (707.1-1414.2 Hz):

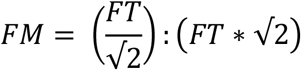

The target and the masking sounds were separated by a changing interval of silence, which was the main independent variable in this experiment. 1000 ms after the offset of the masking noise, a second 10 ms beep, identical to the target (1000 Hz; fade-in and fade-out 2.5 ms; attenuation: −24 Db), was presented after which the participant had 5 s to respond by pressing one of the two response keys whether they have heard the target stimulus using a keyboard. To avoid a possible inter-trial variability in the relative timing of the stimuli, all sounds (i.e. target, mask, response cue) and intervals of silence were created and played within each trial as a single continuous clip of audio.

Half of the participants responded with the right hand button if they could hear the target and with the left if they could not, whereas the response keys were reversed for the other half of participants in order to account for possible lateralization effects in reaction times between the dominant and non-dominant hand responses(71). After a response, or the end of a 5 s response-free period in a case of an omission, there was an 8-12 s pause until the presentation of the next target tone. The inter-trial interval was relatively long in order to allow for drowsiness to develop, which is more difficult to achieve if a task is speeded. Participants were instructed that a target sound will not always be presented, and they did not know that in fact all trials had a target sound, i.e. there were no catch trials during the behavioural staircase experiment.

The first trial always had a detectable target with the masking noise presented after a 400 ms gap, which was considerably longer than the highest threshold among our participants (see below). For the following trials, a simple up-down staircase procedure was applied to determine a required gap between the target and masking sounds(72). That is, the gap was increased if participants were unaware of the target, making the next trial easier, and it was increased if participants could hear the target sound, making the next trial more difficult. The staircase procedure continued until 12 reversals of the change of gap. Until the fourth reversal, the gap duration changed by a factor of 2, and afterwards, until the 12th reversal, it changed by a factor of √2. Threshold was calculated as an arithmetic mean of the gap values during the last 8 reversals. The same staircase procedure was repeated twice, and the mean threshold of these two blocks was used for the main EEG experiment. For the behavioural experiment, the stimuli were created and the staircase procedure was controlled by the Psychoacoustics toolbox(73) running in Matlab (R2011b) on a MacBook Pro.

Staircase procedure revealed a high variability of auditory threshold, i.e. the silent interval between the target and masking sounds required to hear the target, across participants (M=71.03 ms, SD=55.58, Min=6.67 ms, Max=202.4 ms). Nevertheless, despite individual differences, almost identical thresholds were estimated in both blocks of the behavioural experiment (Block 1: M=67.75, SD=52.85; Block 2: M=74.31, SD=61.50; Pearson’s r = 0.89, p<0.000001), confirming the high reliability of the staircase procedure.

### Stimuli during EEG experiment

During the main EEG experiment, the same auditory stimuli were presented as in the behavioural staircase procedure, except that the interval (T) between the target stimulus and the noise, which varied now around the participant’s individual threshold (T_0_). 11 different interval conditions were calculated via:

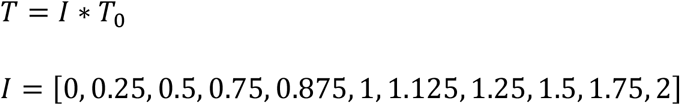

Additionally, a 12^th^ ‘catch trial’ condition in which there was no target stimulus was presented in order to assess the false alarm rate of each participant. The three conditions centred around an individual participant’s threshold (T=[0.875,1,1.125]*T_0_) were four times more likely to be presented in comparison to the other gap conditions and catch trials. All trials were presented in a random order. Stimuli were generated within each trial using Psychoacoustics toolbox(73) and the whole experiment was programmed and controlled using Psychtoolbox version 3(74) running in Matlab (R2011b) on a MacBook Pro. Participants were instructed to respond using the same keys as during the behavioral staircase experiment. They were also explicitly told that they are allowed to fall asleep if they want to. The maximal duration of data collection during EEG experiment was 120 min; however, the experiment was terminated earlier if a participant became too tired or uncomfortable. On average, 501 trials were run per participant (SD=65, Min=361, Max=604).

### EEG acquisition and preprocessing

129-channel EEG data, sampled at 500 Hz and referenced to the vertex, were recorded with the Net Amps 300 amplifier (Electrical Geodesics Inc., Oregon, USA). Conductive gel was applied to each electrode to ensure that the impedance between the scalp and electrodes was kept below 100 kΩ. For the data preprocessing and analyses, EEG channels over forehead, cheeks, and neck were excluded to minimise the influence of eye- and muscle-related noise, retaining 92 channels that covered the scalp. Continuous EEG data were then filtered (high pass: 0.5 Hz; low pass: 40 Hz) and re-referenced to the average of all channels. Afterwards, data were epoched around the onset of the target sound (−4000 ms to 6000 ms) and baseline corrected to the 100 ms preceding the target stimulus. The first 5 trials were deleted for each participant, thereby excluding an approximately 1 min period during which participants often moved in the chair till they settled into a comfortable position. The remaining extremely noisy epochs and EEG channels were manually deleted before running the independent component analysis (ICA) for the final removal of artefacts (such as eye blinks and saccades, heart beat, sweating, etc.). ICA was carried out on relatively clean channels only, and the noisy channels were recalculated by spherical spline interpolation of surrounding channels after deleting ICA components with artefacts. On average, 38 trials (7.6%) were deleted per single participant during EEG pre-processing, and there were on average 463 trials per participant (SD=60, Min=345, Max=584) available for further EEG analyses. Data pre-processing was carried out using EEGlab toolbox for Matlab(75).

### Hori-measure of wakefulness

Given that there is no single widely accepted measure of wakefulness in cognitive experiments, three complementary measures were used to assess the depth of transition from waking to sleep: Hori scoring system, EEG theta/alpha power, and reaction times (RT). A clinical Hori-measure of wakefulness is based on visual scoring of 4 sec segments of continuous EEG recording(48). In Hori system, Stage 1 indicates alpha-dominated relaxation, Stage 9 is marked by complete spindles that coincide with a classical Stage 2 NREM sleep, and other stages in-between reflect the gradual progression of sleep onset and the slowing down of dominating EEG frequencies (see Fig S1A). Sequential analysis of the progression of Hori stages during uninterrupted transition from wakefulness to sleep confirms the validity of the rank order of these stages(76). In addition, several studies showed systematic ERP and EEG spectral power changes throughout the Hori-defined progression of sleep onset(77, 78). For instance, amplitude of mismatch negativity (MMN) decreases from Hori Stages 1 to 3 and then changes its polarity from Hori Stages 4 to 9(77), whereas the peak regions of EEG alpha power move from posterior to anterior regions of scalp with an increasing depth of transition(78). Finally, reaction times as well as the rate of subjective reports of being asleep steadily increase from Hori Stages 1 to 9(48), corroborating the use of this system to measure the depth of wakefulness in humans.

In the present study, Hori stages were visually assessed over 4 sec epochs of pre-stimulus period (see Fig 1A) by two experienced raters, who were blind to the response type (hit, miss, or unresponsive) of each particular trial. For scoring purposes, EEG recordings were low pass filtered (20 Hz), and only 21 EEG channels of the standard 20-10 system were evaluated. In a case of a disagreement (which occurred on 13.2% of trials, usually between any two adjacent Hori stages), the raters discussed the difference until an agreement was achieved. For the EEG analyses, “Hori-awake” trials were defined as Hori Stages 1 and 2, dominated by alpha waves, and trials scored as Hori Stages 4 and 5, dominated by the slower theta waves, were regarded as “Hori-drowsy”.

While being perhaps the most accurate electrophysiological assessment of the level of sleep onset progression(45), the use of Hori system in cognitive neuroscience experiments is however limited by the unpredictable proportion of different stages within an individual participant. In our dataset of 31 participants, distribution of Hori scores ranged from participants having mostly Stage 1 epochs to participants with mostly Stage 5 trials (see Fig S1B), rendering within-participant comparisons between awake and drowsy trials difficult to perform. In such cases, trials of different participants could be grouped by Hori score and merged into a single dataset, following which analysis could carried out at a single trial level, under the assumption that the variation in the relative contribution of individual participants to each Hori score does not contribute significantly to the variance between experimental conditions of interest. An alternative approach would be to carry out analyses at individual participant level following other measures of wakefulness that enable equal division of trials across arousal conditions, such as theta/alpha- or RT-based measures of wakefulness.

### Theta/alpha-measure of wakefulness

Given that Hori Stages 1 to 4 are marked by a decreasing alpha range activity, whereas Stages 4 to 8 have an increasing theta range activity(48) (see Fig S1A), progression of wakefulness can be quantified by the spectral power of respective EEG frequency bands. That is, wakefulness can be defined as a period of time with an increased theta and a decreased alpha band power, or – by combining these two measures – as a period of an increased ratio of theta/alpha (θ/α) power. After calculating theta/alpha ratio for each trial within an individual participant, data can be divided into equal number of relatively “awake” and “drowsy” trials, this way maximizing power of neurocognitive contrasts between these states of consciousness(39, 79).

To apply θ/α measure of wakefulness in the present study, spectral power of EEG frequency oscillations was computed every 4 ms between −2444 ms to 2444 ms in respect to the onset of a target tone using continuous wavelet transform, set from 3 cycles at 3 Hz to 8 cycles at 40 Hz. Theta (4-7 Hz) and alpha (8-12 Hz) power was then averaged individually for each trial from −2000 ms to 0 ms and a theta/alpha ratio was calculated for each electrode. Finally, theta/alpha power was averaged across all electrodes, resulting in a single “sleepiness” value per trial. Trials were then split between the most “θ/α-awake” and the most “θ/α-drowsy” trials, separately for each gap condition. Two data splits were used for each individual participant: a thirds’ split between 33% of the most “awake” and 33% of the most “drowsy” trials (θ/α-33%), and a median split between 50% of the most “awake” and 50% of the most “drowsy” trials (θ/α-50%). Arguably, θ/α-33% measure could avoid uncertain intermediate trials, whereas θ/α-50% measure could maximize on the available number of trials.

### RT-measure of wakefulness

We also used reaction times (RT) as a behavioural measure of the depth of wakefulness at the moment of response execution rather than several seconds before the presentation of stimuli (see Fig 1A). Typically, reaction times are prolonged in a state of low vigilance(80, 81), decreased wakefulness(48, 82), and after sleep deprivation(83–85). Arguably, the speed of processing may become slower in the drowsy state at any stage of information processing: sensory processing, stimuli recognition and categorization, decision-making, motor preparatory processes, and/or execution of motor commands. To narrow down a potential variance of RT-measure of wakefulness to the motor processes, a response cue was presented indicating a time to respond (see Fig 1A). Assuming that a 1000 ms period from the offset of mask to the onset of response cue was sufficient to make a response decision, RT was interpreted as reflecting a period required for a motor plan and its execution.

Importantly, some conflicting evidence from traffic psychology shows no association between RT and sleepiness(86) or sleep debt(87). We thus employed the RT-measure of wakefulness with a caution, treating it as a complementary rather than a central measure in the analyses reported below. Similarly to the θ/α-based split between “awake” and “drowsy” states, trials were separated for each gap condition within each participant into the fastest third of RTs and the slowest third of RTs (RT-33%). In addition, a similar split around the median RT separated data into a half of “awake” trials with the fastest RT and another half of “drowsy” trials with the slowest RTs (RT-50%).

### Agreement between different measures of wakefulness

Where possible, Hori measure of wakefulness was preferred over the θ/α- and RT-measures because its absolute electrophysiological signatures of the transition from wakefulness to sleep can be identified within each participant. When psychophysical analyses required equal number of “awake” and “drowsy” trials within each participant, we considered following θ/α- and RT- measures of wakefulness that allowed such splits of data. As a downside, these measures are relative and even if all trials would be of Hori Stage 1, one third (or half) of them would still get scored as “drowsy”. Thus, aiming to verify the use of θ/α- and RT- data splits, we compared all three measures at an individual as well as at a group level.

First, we carried out correlation analyses between any two measures of wakefulness within each participant, using a raw Hori, θ/α or RT score of all trials within a session, i.e. without splitting them into “awake” and “drowsy” categories (see Fig S1C). Only responsive trials were used for correlations involving RT. Second, we compared correlation coefficients against zero, aiming to assess a consistency of an association between any two measures of wakefulness at a group level (see Fig S1D). Hori and θ/α-measures were positively and significantly correlated for all 31 participants (individual rho ranged from 0.39 to 0.88). Group analysis confirmed a very strong association between these two electrophysiological measures of wakefulness (one sample t test: t(30)=26.16, p<0.000005), confirming that θ/α can be used reliably to assess the level of wakefulness. Contrary to this, correlation coefficients between Hori and RT-measures were less consistent (from −0.15 to 0.41) and significant only in 16 out of 31 participants. Similarly, correlation coefficients between θ/α- and RT-measures ranged from −0.14 to 0.41. While group level analyses showed a significant association between Hori scores and RTs (t(30)=3.75, p=0.001) as well as between θ/α and RTs (t(30)=4.12, p=0.00028), we further treated RT as a complementary rather than a central measure of wakefulness as it was not reliable at a single participant level.

### Behavioural analyses

We aimed to investigate conscious access in awake and drowsy trials by fitting two different models, a sigmoid function and a linear function, and comparing threshold and slope measures. The auditory detection in awake and drowsy trials was first investigated by fitting a sigmoid function to the ratio of hits to misses (constrained from 0 to 1 on the *y* axis) across 11 gap conditions and comparing the threshold and slope measurements between the states of consciousness in each participant separately:

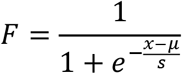

where *F* is the hits ratio, *x* is the gap condition, *µ* is the threshold value (the gap condition at the inflection point), and *s* is inversely proportional to the slope at the threshold. The sigmoid was fit using the Signals Approach Toolbox for Matlab (Spencer Lynn, http://code.google.com/p/satb/).

In order to investigate whether conscious access remained non-linear in the drowsy state, the data were also fitted to a linear function:

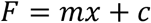

where *F* is the predicted hits ratio, *x* is the gap condition, *m* is the slope, and *c* is the point at which the line crosses the *y* axis.

The goodness of fit was compared for each model in each state of consciousness. As both fitting functions contained the same number of free parameters, 2, the models were compared using the *R^2^* values given by:

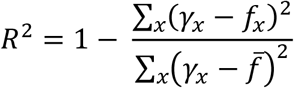

where *γ_x_* is the hits ratio measured for each gap condition *x*, *f_x_* is the predicted hit ratio given by the model, and 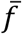 is the mean hits ratio measured over all gap conditions. *R^2^* varies from 0 to 1 with 1 indicating a perfect fit to the data.

### Spatio-temporal ERP clustering

To identify neural signatures of conscious access and its inhibition, auditory ERP dynamics were studied using data-driven spatiotemporal clustering analysis similar to what we previously described(54). Time windows of interest were compared between hits and misses at a single trial level, i.e. by concatenating individual datasets without participant-level averaging. While participant-level averaging is typically performed to contrast independent experimental conditions, a single trial-level analysis is arguably more appropriate when trials of independent conditions fluctuate spontaneously and their count cannot be pre-determined within- and between-participants, such as during rapid transitions between awake and drowsy states (see Fig S1C). To narrow down possible physical variance between the stimuli, only trials corresponding to gap conditions close to participant-wise thresholds (‘-12.5%’, ‘0%’, ‘+12.5%’) were subjected to ERP clustering. Furthermore, ERP clustering was limited to trials of Hori stages 1-5, as ERP topography tended to change in Hori stages ≥6.

Using functions of FieldTrip toolbox(88, 89), we compared corresponding spatiotemporal points in individual awake and drowsy trials with an independent samples t test. Although this step was parametric, FieldTrip used a nonparametric clustering method(90) to address the multiple comparisons problem. t values of adjacent spatiotemporal points with p<0.05 were clustered together by summating them, and the largest such cluster was retained. A minimum of two neighbouring electrodes had to pass this threshold to form a cluster, with neighbourhood defined as other electrodes within a 4 cm radius. This whole procedure, i.e., calculation of t values at each spatiotemporal point followed by clustering of adjacent t values, was repeated 1000 times, with recombination and randomized resampling before each repetition. This Monte Carlo method generated a nonparametric estimate of the p value representing the statistical significance of the originally identified cluster. The cluster-level t value was calculated as the sum of the individual t values at the points within the cluster. Spatiotemporal clustering was always carried out within a restricted time window around the ERP peak of interest (P50, N100, P200, or P300).

First, a butterfly plot of trials locked to the onset of mask and taken together from all participants (N=31) revealed 3 prominent peaks at 92 ms, 196 ms, and 528 ms after the onset of mask (see Fig S3C). Based on their topography and latency, the peaks were identified as N100, P200 and P300 potentials (see Fig S3C). Afterwards, hits (N=3442) and misses (N=3277) were subjected to spatiotemporal ERP clustering around the time window of each individual peak, namely a ± 50 ms time window around the 92 ms peak for N100 potentials, a ± 50 ms time window around the 196 ms peak for P200 potentials, and a ± 250 ms time window around the 528 ms peak for P300 potentials (Fig S3D-F).

To isolate relatively early responses to the target, EEG data were selected from 10 participants who showed the highest thresholds in the behavioural staircase experiment (M=139.66 ms, SD=35.86 ms, range: 83.55-202.4 ms). Given that for these participants the masking noise was presented on average 140 ms after the onset of the target, and assuming that the earliest responses to the mask would arise around > 30 ms after its onset (see Fig 2C), we expected that potentials in a time window < 170 ms will be driven by the target sound. When trials of these 10 participants were grouped together, a butterfly plot of hits and misses revealed two prominent peaks at 50 ms and 116 ms (see Fig S3A). Trials were then separated between hits (N=1527) and misses (N=842), and data-driven clustering at a single-trial level was carried out separately in a ± 25 ms time window around the P50 peak, and in a ± 50 ms time window around the N100 peak (see Fig S3B).

### ERP conditioning and normalization

Following spatio-temporal clustering, neuronal responses to the mask between awake and drowsy trials were compared across 11 gap conditions. For this, awake and drowsy datasets were created by concatenating corresponding trials of all 31 participants. Next, trial counts were matched between awake and drowsy datasets by deleting randomly selected trials from the awake condition. The following numbers of trials were obtained across 11 gap conditions: 193, 191, 180, 164, 721, 726, 739, 188, 170, 194, and 203 trials in each state of consciousness. To contrast neural markers of hits and misses in awake and drowsy states, four datasets were created to study target-evoked N100 by appending trials of 10 participants who had the highest target detection threshold: awake misses (N=363), awake hits (N=931), drowsy misses (N=319) and drowsy hits (N=320). All four conditions were matched to have 319 trials each by deleting randomly selected trials from the awake misses, awake hits and drowsy hits conditions. Similarly, four single-trial datasets were created to study mask-evoked N100, P200 and P300 components, this time appending trials of all 31 participants: awake misses (N=1593), awake hits (N=1759), drowsy misses (N=793) and drowsy hits (N=803). All four conditions were then matched to have 793 trials each by deleting randomly selected trials from the awake misses, awake hits and drowsy hits conditions.

To normalise within-trial variance, raw voltage of each individual trial was transformed to z-scores using the mean and standard deviation of the baseline period (−100 to 0 ms). As in the preceding spatiotemporal ERP clustering, 0 ms indicated the onset of a target tone when target-evoked N100 was studied, and the onset of a mask when mask-evoked ERP components were studied. Next, to correct for between-trial variance introduced by inter-individual EEG differences in the drowsy state of consciousness, cross-trial weighting of variance was performed. Namely, each data point along a time axis of a single trial was multiplied by the absolute value of the mean divided by the standard deviation of the same time point across all trials and all conditions:

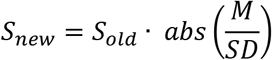

where *S_new_* is a new weighted single time sample of an individual trial, *S_old_* is an original value of the same time sample of an individual trial, *M* is the mean of all trials across all conditions at the same time sample, and *SD* is the standard deviation of all trials across all conditions at the same time sample. Cross-trial weighting reduces amplitude of time points that show relatively high inter-trial variance (induced activity) and amplifies signal with relatively high inter-trial consistency (evoked activity). Importantly, ERP cross-trial weighting is sensitive to the number of trials included, and thus all conditions in each ERP analysis were matched by a trial count. Both standardization and cross-trial weighting were carried out separately for each electrode, i.e. there was no spatial correction implemented in the analysis. For the EEG slope analysis, cross-trial weighting was carried out by calculating joint M and SD over awake and drowsy trials separately for each gap condition, which had physically different stimuli. For the ERP analysis of States x Access interaction and ERP source modelling, cross-trial weighting was carried out by calculating joint M and SD over trials in all four conditions included in the analysis, i.e. awake misses, awake hits, drowsy misses, and drowsy hits.

### EEG slope analysis

To study the sensory-central gating and its modulation by the state of consciousness, EEG epochs locked to the onset of mask were divided between Hori-defined awake (stages 1-2) and drowsy (stages 4-5) conditions. Next, trials were averaged separately for the 11 gap conditions, and this was repeated for the awake and the drowsy trials. Mean amplitude of mask-evoked ERP clusters in N100, P200, and P300 time windows was then averaged separately for each gap condition and both states of consciousness (see below). This reduced data to six two-dimensional matrices of 92 electrodes x 11 gap conditions, i.e. one matrix for each state (awake, drowsy) and time window (N100, P200, P300). Next, for each electrode, a linear function was fitted to 11 amplitude values (one per gap condition), and a slope and a coefficient of determination (R^2^) were calculated, using the same functions as for behavioural analysis. The same procedure was repeated for all six matrices of awake and drowsy datasets at N100, P200 and P300 time windows. Finally, the obtained slope and R^2^ raw values were plotted as topographical maps, and their absolute values were used for statistical analyses (see Fig 3 and Fig S6).

To visualize changing slope across 11 gap conditions, the identified ERP markers of conscious access were analysed using a measure of global field power (GFP)(Lehmann & Skrandies, 1980) within a time window of interest. GFP was calculated as a standard deviation of raw voltage values at a single time point of individually averaged waveforms across all 92 electrodes. This way, GFP reduces spatial dimension of EEG data to a single value as a parametric summary of a momentary strength of topographical EEG map. Given that GFP was calculated from weighted EEG data, we refer to the obtained measure as weighted GFP (wGFP).

### ERP analysis of States x Access interaction

For the analysis of Access x States interactions of the identified N100, P200 and P300 spatio-temporal clusters (see above), cluster peaks were reassessed by plotting a waveform of averaged cluster electrodes in gap conditions ‘-12.5%’, ‘0%’ and ‘+12.5%’, and detecting a peak within a significant cluster time window. While doing this, four conditions of interest – awake misses, awake hits, drowsy misses, and drowsy hits – were matched by the trial count and then averaged. Waveform peaks were searched within significant time windows of originally identified clusters, and the following time windows were used to calculate mean amplitude as close around the peak as possible without going beyond a cluster significant time windows: 50 ms for N100, 80 ms for P200 and 120 ms for P300. This way, the following time windows were identified and used to calculate mean amplitude: 118-166 ms for the positive target-evoked N100 cluster (mean amplitude constrained by cluster significant window of 48 ms), 76-126 ms for the negative mask-evoked N100 cluster, 166-246 ms for the positive mask-evoked P200 cluster, and 362-482 ms for the positive mask-evoked P300 cluster. The same cluster time windows were also used for the ERP slope analysis (see above) as well as the ERP source modelling (see below). Weighted amplitude of a respective ERP component (N100, P200, or P300) was averaged across cluster electrodes and identified time samples for the statistical analyses.

### EEG source modelling

Weighted minimum norm estimation (wMNE) of single trial sources was carried out to study whether temporal and fronto-parietal sources contribute to the States x Access interaction. Source modelling was performed with Brainstorm(91), which is documented and freely available for download online under the GNU general public license (http://neuroimage.usc.edu/brainstorm). A head model was computed using OpenMEEG BEM forward modelling(92, 93), with the following BEM layers and conductivities: scalp: 1082 vertices – 1; skull: 642 vertices – 0.0125; brain: 642 vertices – 1. The canonical Colin27 was used as a structural magnetic resonance imaging (MRI) template to create these surfaces. Noise covariance computed across all channel pairs was incorporated in the wMNE algorithm. To draw scouts of the hypothesized sources in the expected regions of interest (ROIs), absolute values of source activations within each scout were averaged across cluster peak time window that was determined during ERP analyses (see above), trials and all four conditions, i.e. awake misses, awake hits, drowsy misses, and drowsy hits (793 trials per condition). N100 and P200 time windows guided identification of joint temporal lobe source, and P300 time window guided identification of frontal and parietal sources. The size of the scouts was fixed at 85 cm^2^, centred on the cortical peak of the average activation. Source activation of each trial and condition within selected scouts was subjected to statistical analysis.

### Statistics

Statistical analyses were carried out using IBM SPSS Statistics and Matlab R2014a. When possible, parametric tests were used. When Levene’s test indicated different amounts of variability between scores of two conditions, equal variances were not assumed when running an independent-samples t test. Cohen’s d was calculated to assess effect size of pairwise comparisons, using pooled variance. In cases when data were ordinal, non-parametric tests were used.

Regarding specific contrast conditions: **(i)** For within-participant comparisons of different measures of wakefulness (Hori, θ/α, RT), the Spearman rank order correlation test was used, as Hori stages are ordinal. The obtained correlation coefficients were subjected to group level analysis with a one-sample t test. **(ii)** Unequal spread of the rate of hits across 11 gap conditions was tested using one-way repeated measures ANOVA. **(iii)** To compare threshold and slope estimates between awake and drowsy trials, a separate paired sample t test was carried out for each split of data, i.e. θ/α-33%, θ/α-50%, RT-33%, and RT-50%. **(iv)** To compare R^2^ of sigmoid versus linear fits of the hits’ rate across 11 gap conditions between awake and drowsy states of consciousness, repeated measures ANOVA was carried out separately for the θ/α-33% and RT-33% based splits with Models (linear, sigmoid) and States (awake, drowsy) as independent factors. Pairwise comparisons were carried out using paired samples t tests. **(v)** To compare a difference score of the rate of hits between the easiest and the most difficult gap conditions across Hori Stages 1-6, a one way between samples ANOVA with a linear contrast was carried out with Hori stages as an independent factor and the difference score of hits between the easiest and most difficult gap conditions as a dependent factor. In this trend analysis, a weighted linear term was used to compensate for unequal size of observations across Hori stages. **(vi)** Association between slope and threshold estimates in awake and drowsy trials was evaluated using Pearson correlation, which was carried out separately for the θ/α-33%, θ/α-50%, RT-33%, and RT-50% splits of the data. **(vii)** To evaluate state-induced modulation of ERP-clusters of conscious access at a single trial-level, two-way between-trials univariate ANOVA was carried out with States (awake, drowsy) and Access (miss, hit) as fixed factors, separately for N100, P200, and P300 components. Independent samples t tests were used for follow up analyses of significant main effects and interactions. **(viii)** To evaluate state-induced modulation of the sources of mask-evoked ERP-clusters at a single trial-level, two-way between-trials univariate ANOVA was carried out with States (awake, drowsy) and Access (miss, hit) as fixed factors and the amplitude of the weighted current density averaged across 6 scouts (left frontal, right frontal, left parietal, right parietal, left temporal, right temporal) as the dependent variable, separately for N100, P200, and P300 components. In a case of a significant main effect of Access, planned independent samples t tests were carried out to contrast hits and misses in awake and drowsy states in frontal, parietal and temporal sources, aiming to identify sources with the higher amplitude in misses than hits. **(ix)** To compare the slope of a linear fit across the mean ERP amplitude of 11 gap conditions between awake and drowsy trials in 92 electrodes, repeated measures ANOVA was carried out with ERP component (N100, P200) and States (awake, drowsy) as fixed factors. Identical test was carried out to analyse R^2^ of these fits. Paired sample t tests were used for follow up analyses of significant main effects and interactions. **(x)** To compare ordinal Hori scores of sleep progression between the excluded (N=25) and the remaining (N=31) participants, the mode of Hori scores was first calculated for each participant across the whole EEG session, and these scores were then compared between the groups using non-parametric Mann-Whitney U test. **(xi)** Pearson χ^2^ test was used to test if excluded participants were more likely to be male or female compared to the remaining participants. **(xii)** Pearson correlation was used to assess the strength of association between detection threshold estimates in Block 1 and Block 2 of the behavioural staircase procedure.

## Acknowledgement

This research was supported by a Wellcome Trust Biomedical Research Fellowship WT093811MA (TAB) and an IRSES EU Exchange (TAB).

## SUPPLEMENTARY RESULTS

### Neural markers of conscious access and its suppression

Given that both sensory and executive stages of processing may index conscious access(5, 15), and that the target and the masking stimuli were presented serially, we considered that reported awareness of a target might be associated with an increase of target-evoked event-related potentials (ERP), whereas suppression of conscious access might be marked by the higher amplitude of mask-evoked N100, P200 and P300 responses. Thus, we first contrasted respective potentials locked to the onset of mask by carrying data-driven spatiotemporal clustering of hits and misses.

In this analysis, all data were included with no split between awake and drowsy trials. We found a significantly more negative fronto-central N100 cluster from 42 ms to 142 ms (see Fig S3 D) and a significantly more positive occipital cluster from 42 ms to 134 ms in misses compared hits. Likewise, clustering of P200 potentials revealed that misses were associated with a more positive central cluster (150-246 ms; Fig S3 E) and a more negative frontal cluster (146-246 ms) than hits. Finally, misses were associated with more positive parieto-occipital P300 cluster (290-482 ms; Fig S3 F) and more negative frontal cluster (278-530 ms) than hits. In other words, when participants reported awareness of the target sound, the amplitude of early (N100), middle (P200) and late (P300) responses to the mask was relatively low. Conversely, when conscious access was suppressed (missing the target when present), the mask evoked relatively higher amplitude of neural responses.

We next investigated if ERP to the target sound would also distinguish hits and misses, this time revealing neural signatures of conscious access rather than its suppression. Despite the constrains of the design, gap events were long-enough to assess target-specific N100 responses in a subgroup of participants (N=10). We found a significantly more negative central N100 cluster of electrodes from 118 ms to 166 ms (see Fig S3 B) and a more positive occipital cluster of electrodes from 112 ms to 166 ms in hits compared to misses. That is, hits or “aware” trials had higher amplitude of N100 response to the target tone, pointing to a relatively early sensory involvement of auditory conscious access. Given that the target and the masking sounds were relatively adjacent in time, i.e. the mean gap of silence between them was 71 ms (range: 7-202 ms), evoked potentials to the target and mask largely overlapped, occluding the analysis of P200 and P300 responses specific to the target. No reliable ERP differences between hits and misses were found within the P50 time window.

In summary, spatiotemporal clustering of ERP differences between hits and misses revealed a cascade of electrophysiological markers at different depths of processing that distinguished conscious and unconscious trials. In particular, conscious access was associated with enhanced N100 response to the target, whereas the suppression of conscious access was associated with enhanced amplitude of potentials evoked by the mask (N100, P200, P300).

## SUPPLEMENTARY TABLE

**Table 1.**
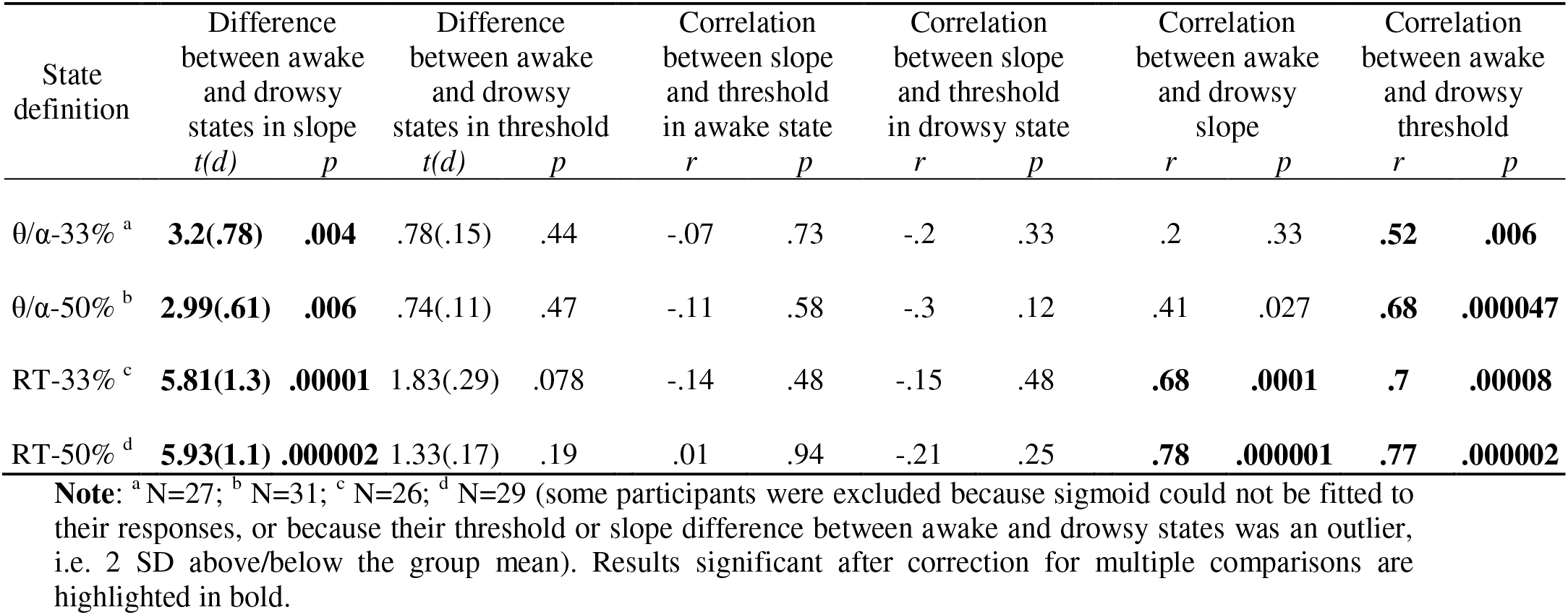
Slope and threshold estimates in awake and drowsy states: statistics

## SUPPLEMENTARY FIGURES

**Figure S1.**
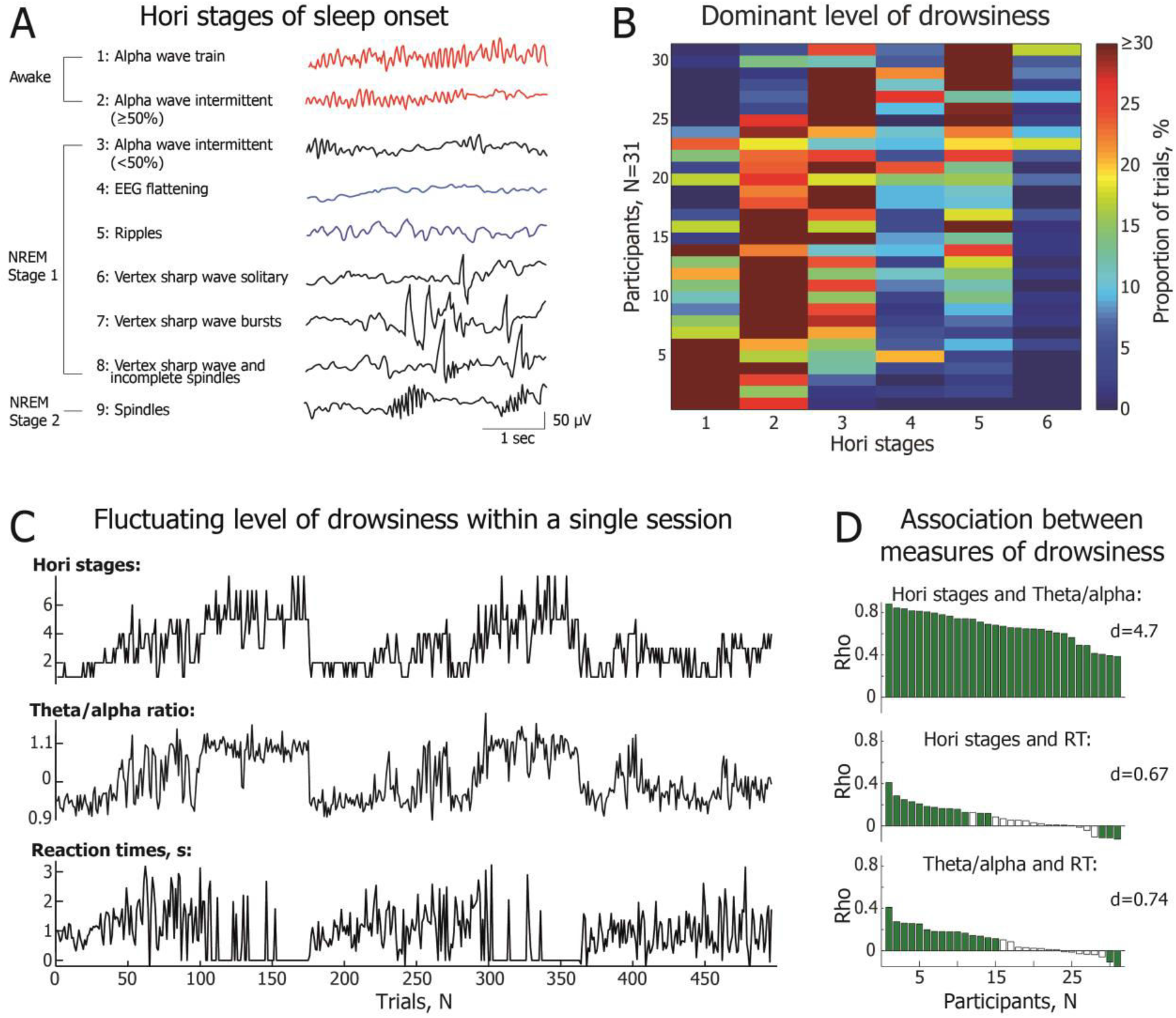
EEG and RT measurements of wakefulness. **(A)** EEG examples and brief definitions of 9 Hori stages of sleep onset progression from the relaxed wakefulness to the non-rapid eye movement (NREM) Stage 2 sleep (modified with permission from(45)). Hori stages 1-2 defined awake trials (marked in red), whereas Hori stages 4-5 defined drowsy trials (marked in blue). **(B)** Percentage of trials scored as different Hori stages within each participant whose data were used for the analyses of conscious access. Datasets are sorted here from the most alert participants 1-6 with the dominance of Hori stage 1 to the drowsiest participants 30 and 31 with the dominance of Hori stage 5. There were relatively few epochs of Hori stage 4 (EEG flattening is known to last for a relatively short period of time(94)) and Hori stages 6 and above. **(C)** A representative dataset of one participant with good agreement between the three measures of wakefulness across 500 consecutive trials. The top subplot indicates Hori stages, the middle subplot depicts θ/α ratio, and the bottom subplot shows fluctuation of RT. An RT of 0 indicates an unresponsive trial, whereas a negative RT indicates a premature response that took place before the response cue. **(D)** Correlations between different measures of wakefulness: Hori stages vs. θ/α power (top), Hori stages vs. RT (middle), and θ/α power vs. RT (bottom). Bars represent intra-individual Spearman’s rank order correlation coefficients between the respective measures of wakefulness for 31 participants, sorted from the most positive to the least positive correlation coefficients. Green bars represent participants with a significant intra-individual correlation, whereas white bars represent participant who did not show a significant correlation between two respective measures of wakefulness. Group level analyses of the correlation coefficients were carried out using one-sample t tests with the following results: Hori stages vs. θ/α power (t(30)=26.16, p<0.000005, d=4.7), Hori stages vs. RT (t(30)=3.75, p=0.0008, d=0.67), and θ/α power vs. RT (t(30)=4.12, p=0.0003, d=0.74). While some individuals showed a consistent pattern of state-fluctuation across all three measures of wakefulness, a very high convergence was observed only between Hori and θ/α measures of wakefulness.

**Figure S2.**
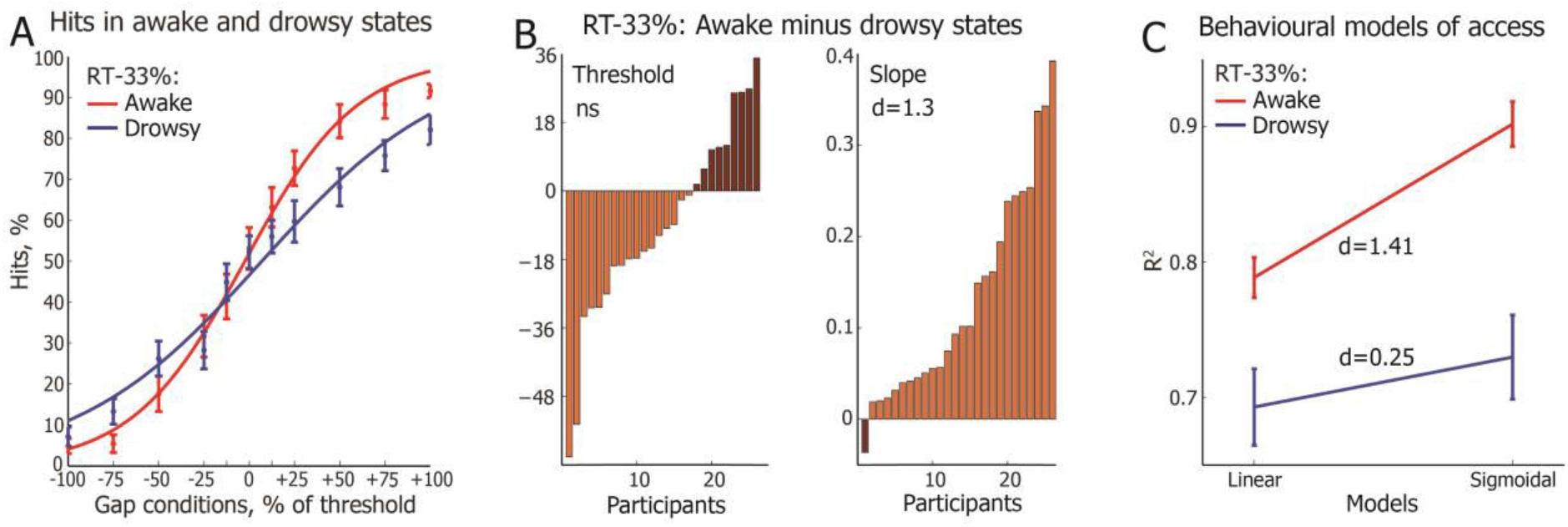
Slope and threshold estimates in RT-defined relaxation and wakefulness. **(A)** Sigmoidal functions fitted to the awake (red) and drowsy (blue) hits, following RT-33% definitions of the states of consciousness. The trials were first averaged within each participant and then across all participants. The error bars indicate the standard error of mean (SEM), calculated across all participants. **(B)** Threshold and slope estimates in the drowsy state subtracted from those in the awake state. Individual participants (N=29), represented as bars, are sorted from the one with the largest increase of threshold (light brown) to the one with the largest decrease (dark brown) of threshold in wakefulness (left), and from the one with the largest increase of slope (light brown) to the one with the largest decrease of slope (dark brown) in wakefulness (right). Data split in this plot is based on the RT-33% definition of the awake and drowsy states. **(C)** Coefficient of determination (R^2^) of linear and sigmoidal models of the proportion of hits as a function of gap in the awake and drowsy trials as defined by RT-33%. The sigmoidal model had a higher R^2^ than the linear model in the drowsy (t(25)=3.86, p=0.0007, d=0.25) and even more so in the awake (t(25)=7.35, p=1.1E-7, d=1.41) state. The error bars indicate the SEM.

**Figure S3.**
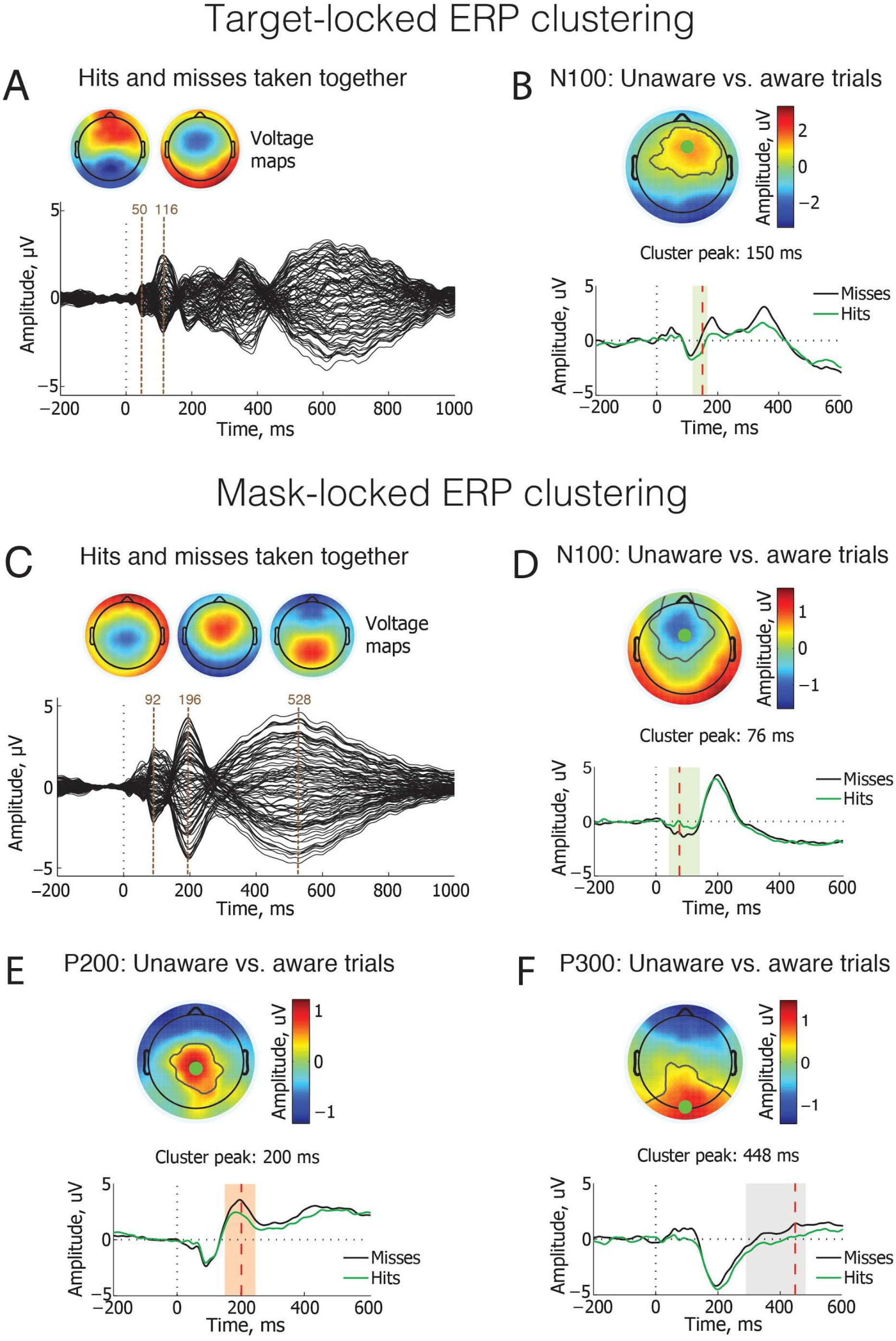
ERP markers of the suppression of conscious access. **(A)** A butterfly plot of ERP waveforms locked to the onset of target, baseline corrected from −100 ms to 0 ms. Topographical voltage maps depict spatial distribution of the butterfly peaks at 50 ms and 116 ms (brown verticals), revealing typical distribution of P50 and N100 auditory potentials. A relatively late 116 ms latency of the N100 peak likely depended on a weak intensity of the target sound (single frequency, short duration, low amplitude), as the N100 latency increases with a decreasing intensity of auditory stimuli(95, 96). These data were taken from 10 participants with the highest threshold of target detection, i.e. the longest period of silence between the target and the masking tones. While the first two peaks reflect neuronal processing of target, the mask, timing of which varied within and between participants, likely contributed to the peaks above 170 ms. **(B)** Spatiotemporal clustering of ERPs locked to the target sound. Hits (green waveform) and misses (black waveform) were compared within N100 time window for 10 participants with the highest auditory detection threshold. Green shaded vertical bar behind the waveforms represents time window of a significant difference between hits and misses (cluster t=2373.1, p=0.001). These waveforms are taken from the fronto-central electrode with the largest difference between hits and misses, marked as a green dot in the topographical voltage map. The black contours within the map reveal which electrodes showed a significant difference, i.e. the more negative amplitude in hits than misses, and formed a spatiotemporal cluster. Red vertical line at 150 ms in the waveform reveals the peak time of the largest difference between hits and misses. The topographical voltage map depicts the very same time point of the most significant difference between conditions. **(C)** A butterfly plot of ERP waveforms locked to the onset of mask, baseline corrected from −100 ms to 0 ms. Topographical voltage maps depict spatial distribution of the butterfly peaks at 92 ms, 196 ms, and 528 ms (brown verticals), revealing typical distribution of N100, P200 and P300 auditory potentials. Data are taken from all 31 participants. **(D-F)** Spatiotemporal clustering of ERPs (hits vs. misses) locked to the masking sound within N100, P200 and P300 time windows. Misses were associated with significantly more negative amplitude of **(D)** N100 peak (cluster t=3674.8, p=0.001), and more positive amplitude of **(E)** P200 peak (cluster t=3327.3, p=0.001) and **(F)** P300 peak (cluster t=5534.7, p=0.004) compared to the hits. In **(D-F)**, grey shaded vertical bars behind the waveforms represent time window of a significant difference between hits (green waveform) and misses (black waveform): green for the N100 cluster, orange for the P200 cluster, and grey for the P300 cluster. The waveforms are taken from the electrode with the largest difference between hits and misses, marked as a green dot in the topographical voltage map. The black contours within the map reveal which electrodes showed a significant difference, i.e. the higher amplitude in misses than hits, and formed a spatiotemporal cluster. Red vertical lines in the waveforms reveal the peak time of the largest difference between hits and misses. The topographical voltage maps depicts the very same time point of the most significant difference between conditions.

**Figure S4.**
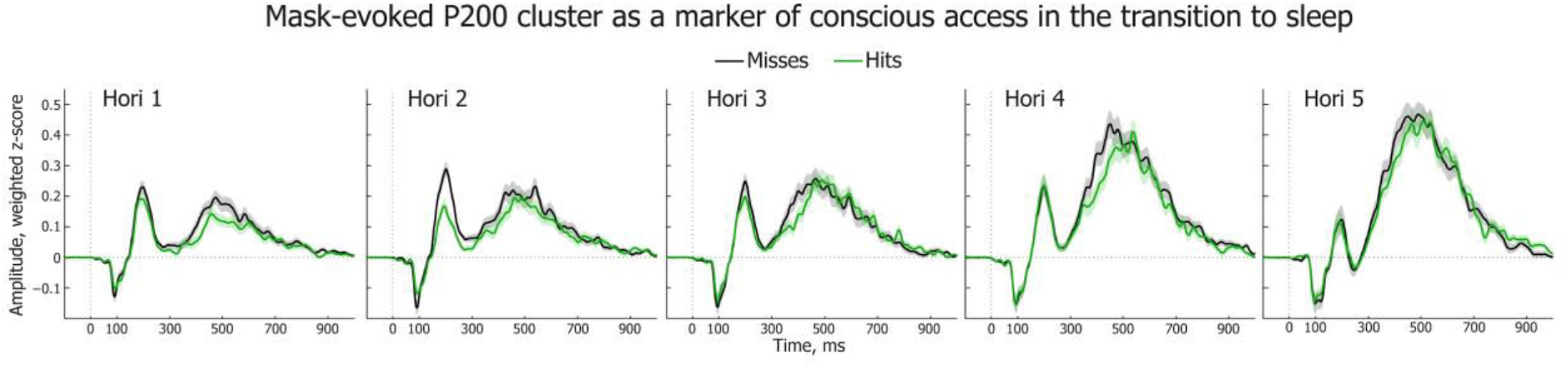
Mask-evoked P200 cluster across five Hori stages of wakefulness. Please note non-linear dynamics of mask-evoked P200 cluster for misses (green) and hits (black) across Hori Stages 1-5, with the suppression of conscious access peaking in Hori Stage 2 (relaxed awake state). Error bars in this figure indicate SEM.

**Figure S5.**
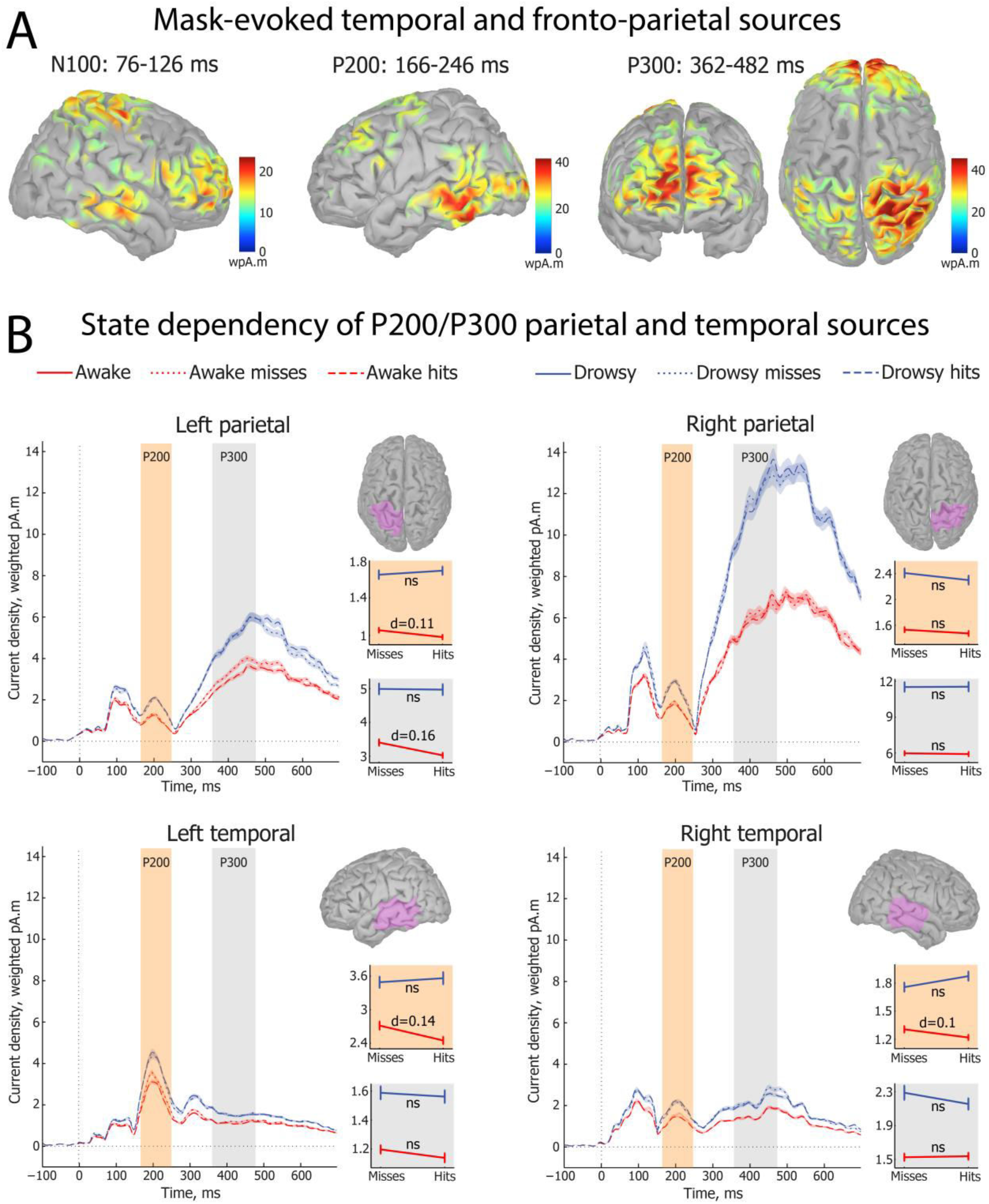
Parietal and temporal cortical sources of mask-evoked ERP clusters. **(A)** Weighted minimum norm estimation (wMNE) of single trial sources averaged across all four conditions in the N100, P200 and P300 cluster time windows. N100 and P200 clusters were used to identify temporal lobe sources and P300 cluster revealed fronto-parietal sources of mask-evoked responses. The sources were modelled using weighted EEG data dipole current density. **(B)** The planned t test comparisons of parietal and temporal sources activations between hits and misses in awake and drowsy states in P200 (orange) and P300 (grey) time windows. Waveforms indicate time courses of source activations, whereas statistical plots depict reliability level of the planned comparisons within respective time windows. Error bars indicate SEM. In the P200 time window, awake misses had higher amplitude than awake hits in the left parietal (t(1584)=2.11, p=0.035, d=0.11) sources, left temporal (t(1560.6)=2.73, p=0.0063, d=0.14) and the right temporal (t(1520.6)=2, p=0.045, d=0.1) sources. In the P300 time window, awake misses had higher amplitude than awake hits in the left parietal (t(1529.7)=3.15, p=0.0017, d=0.16) sources. Drowsy misses showed no higher amplitude than drowsy hits in the sources assessed (only the P300 time in the left frontal source shown in figure X in the main text).

**Figure S6.**
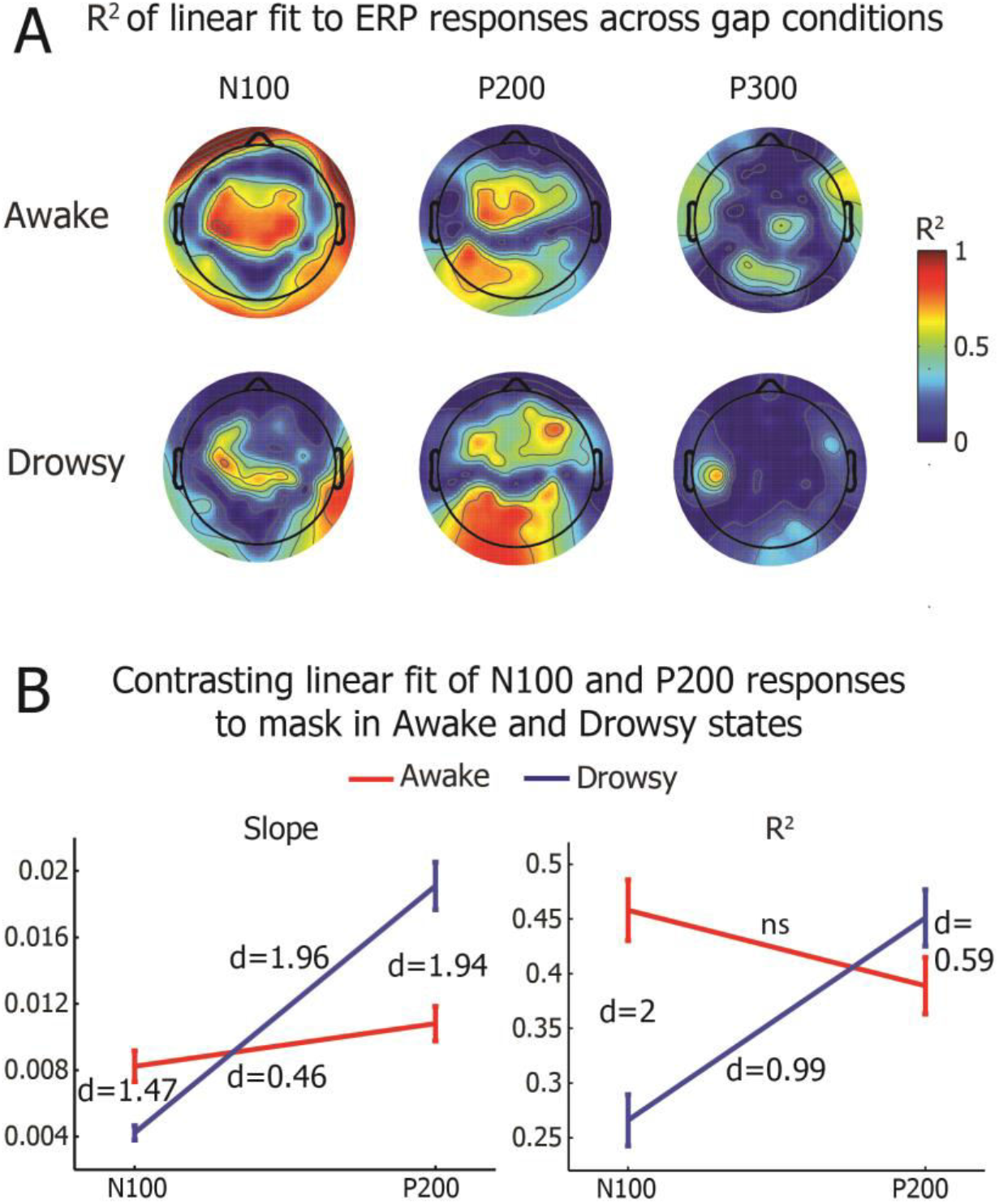
Interaction between ERP slope and states of consciousness. **(A)** Heat maps represent R^2^ values of a linear function, which was fitted to the mean amplitude of mask-locked N100 (left), P200 (middle) and P300 (right) ERP clusters across 11 gap conditions. The R^2^ of a fitted function was first calculated for each electrode and the obtained values were interpolated in the heat maps. The top line represents fit of the awake trials (Hori 1-2), whereas the bottom line depicts fit of the drowsy trials (Hori 4-5). **(B)** Statistical interactions of the heat maps shown in Fig 3 and Fig S6 A. Repeated measures ANOVA was carried out on 92 slope (on the left) or R^2^ values (on the right), representing each electrode, with the State (Awake, Drowsy) and Component (N100, P200) as independent factors. The plots show the mean slope or R^2^ of all electrodes, and the error bars represent SEM. Statistical results of the slope values are fully described in the main text. Analysis of R^2^ maps yielded significant main effect of the States (F(1,91)=17.56, p=0.00006) but not of the ERP components (F(1,91)=2.67, p=0.11), and a significant ERP components x States interaction (F(1,91)=79.64, p<0.000005). Coefficient of determination differed between N100 and P200 components in drowsy (t(91)=4.7, p=0.00001, d=0.99) but not in awake (t(91)=1.8, p=0.076, d=0.38) states of consciousness. N100 had higher R^2^ in awake than in drowsy state (t(91)=9.52, p<0.000005, d=2). Contrary to this, P200 had higher R^2^ in drowsy than in awake state (t(91)=2.83, p=0.0058, d=0.59).

